# Understanding Tissue-specific Gene Regulation

**DOI:** 10.1101/110601

**Authors:** Abhijeet R. Sonawane, John Platig, Maud Fagny, Cho-Yi Chen, Joseph N. Paulson, Camila M. Lopes-Ramos, Dawn L. DeMeo, John Quackenbush, Kimberly Glass, Marieke L. Kuijjer

**Affiliations:** Channing Division of Network Medicine, Department of Medicine, Brigham and Women’s Hospital, Harvard Medical School, Boston, MA; Department of Biostatistics and Computational Biology, Dana-Farber Cancer Institute, Boston, MA; Department of Biostatistics, Harvard T.H. Chan School of Public Health, Boston, MA; Department of Cancer Biology, Dana-Farber Cancer Institute, Boston, MA

## Abstract

Although all human tissues carry out common processes, tissues are distinguished by gene expression patterns, implying that distinct regulatory programs control tissue-specificity. In this study, we investigate gene expression and regulation across 38 tissues profiled in the Genotype-Tissue Expression project. We find that network edges (transcription factor to target gene connections) have higher tissue-specificity than network nodes (genes) and that regulating nodes (transcription factors) are less likely to be expressed in a tissue-specific manner as compared to their targets (genes). Gene set enrichment analysis of network targeting also indicates that regulation of tissue-specific function is largely independent of transcription factor expression. In addition, tissue-specific genes are *not* highly targeted in their corresponding tissue-network. However, they do assume bottleneck positions due to variability in transcription factor targeting and the influence of non-canonical regulatory interactions. These results suggest that tissue-specificity is driven by context-dependent regulatory paths, providing transcriptional control of tissue-specific processes.

## 1. INTRODUCTION

Although all human cells carry out common processes that are essential for survival, in the physical context of their tissue-environment, they also exhibit unique functions that help define their phenotype. These common and tissue-specific processes are ultimately controlled by gene regulatory networks that alter which genes are expressed and control the extent of that expression. While tissue-specificity is often described based on gene expression levels, we recognize that, by themselves, individual genes, or even sets of genes, cannot adequately capture the variety of processes that distinguish different tissues. Rather, biological function requires the combinatorial involvement of multiple regulatory elements, primarily transcription factors (TFs), that work together and with other genetic and environmental factors to mediate the transcription of genes and their protein products [1, 2].

Gene regulatory network modeling provides a mathematical framework that can summarize the complex interactions between transcription factors, genes, and gene products [3–6]. Despite the complexity of the regulatory process, the most widely used network modeling methods are based on pairwise gene co-expression information [7–10]. While these correlation-based networks may provide some biological insight concerning the associations between both tissue-specific and other genes [11, 12], they do not explicitly model key elements of the gene regulatory process.

PANDA (Passing Attributes between Networks for Data Assimilation) is an integrative gene regulatory network inference method that models the complexity of the regulatory process, including interactions between transcription factors and their targets [13]. PANDA uses a message passing approach to optimize an initial network between transcription factors and target genes by integrating it with gene co-expression and protein-protein interaction information. In contrast to other network approaches, PANDA does not directly incorporate co-expression information between regulators and targets. Instead, edges in PANDA-predicted networks reflect the overall consistency between a transcription factor’s canonical regulatory profile and its target genes’ co-expression patterns. A number of studies have shown that analyzing the structure of the regulatory networks estimated by PANDA can help elucidate the regulatory context of genes and transcription factors and provide insight in the associated biological processes [14–17].

The transcriptomic data produced by the Genotype-Tissue Expression (GTEx) consortium [18] provide us with an unprecedented opportunity to investigate the complex regulatory patterns important for maintaining the diverse functional activity of genes across different tissues in the human body [19, 20]. These data include high-throughput RNA sequencing (RNA-Seq) information from 551 research subjects, sampled from 51 post-mortem body sites and cell lines derived from two tissue types.

In this study, we apply PANDA to infer gene regulatory networks for thirty-eight different tissues by integrating GTEx RNA-Seq data with a canonical set of transcription factor to target gene edges (based on a motif scan of proximal promoter regions) and protein-protein interactions. We then use these tissue-networks to identify tissue-specific regulatory interactions, to study the tissue-specific regulatory context of biological function, and to understand how tissue-specificity manifests itself within the global regulatory framework. By studying the structure of these networks and comparing them between fitissues, we are able to gain several important insights into tissue-specific gene regulation. Our overall approach is summarized in Figure 1.

**Figure 1:**
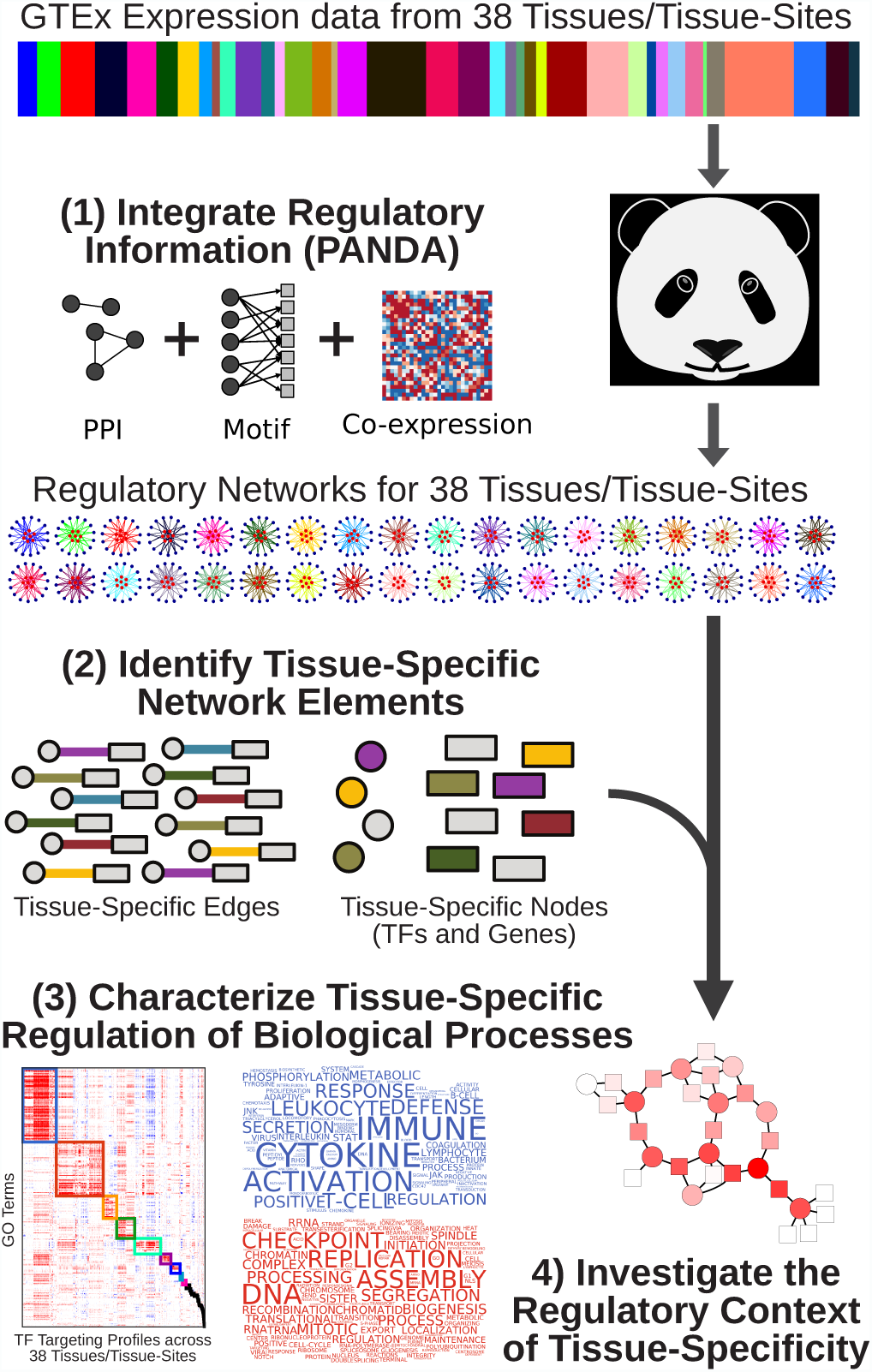
Schematic overview of our approach to characterize tissue-specific gene regulation using the GTEx expression data. We started with gene expression for 9, 435 samples across 38 tissues; the relative sample size of each of the 38 tissues in the GTEx expression data is shown in the color bar. We used PANDA to integrate this information with protein-protein interaction and transcription factor target information (based on a genome-wide motif scan that included 644 transcription factor motifs). This produced 38 inferred gene regulatory networks, one for each tissue. We identified tissuespecific genes, transcription factors, and regulatory network edges and analyzed their properties within and across tissues.

## 2. RESULTS

### 2.1 Identifying Tissue-Specific Network Edges

We started by reconstructing genome-wide regulatory networks for each human tissue. We downloaded GTEx RNA-Seq data from dbGaP (phs000424.v6.p1, 2015-10-05 release). The RNA-Seq data were preprocessed and then normalized in a sparse-aware manner [21] so as to retain genes that are expressed in only a single or small number of tissues. After filtering and quality control, our RNA-Seq data included expression information for 30, 243 genes measured across 9, 435 samples and 38 distinct tissues (Supplemental Materials and Methods). For each tissue, we used PANDA to integrate gene-gene coexpression information from this data set with an initial regulatory network based on a genome-wide motif scan of 644 transcription factors [22] and pairwise transcription factor protein-protein interactions (PPI) from StringDb v10 [23] (Figure 1 and Supplemental Materials and Methods). This resulted in 38 reconstructed gene regulatory networks, one for each tissue. Data and code to reconstruct these networks can be found at http://goo.gl/DE1abB. The reconstructed networks are also available on Zenodo [24].

We used these reconstructed networks to identify tissue-specific network edges. Each PANDA network contains scores (or weights) for every possible transcription factor to gene interaction. We compared the weight of each edge in a particular tissue to the median and interquartile range of that edge’s weight across all 38 tissues. Edges identified as “outliers” in a particular tissue (those with a weight in that tissue greater than the median plus two times the interquartile range of the weight across all tissues) were designated as “tissue-specific.” Using this metric we identified just over five million tissue-specific edges (26.1% of all possible edges, Supplemental Figure S1A, Supplemental Table 1). We compared these tissue-specific edges with other available sources of tissue-specific network relationships [25] (see Supplemental Figure S2 and Supplemental Material and Methods) and found minimal overlap, indicating that these edges may highlight important regulatory interactions that have not been previously explored.

Figure 2A shows the number of edges identified as specific in each of the 38 tissues, colored based on their “multiplicity,” or the number of tissues in which an edge is identified as specific. We found that the majority of tissue-specific edges (65.7%) have a multiplicity of one, meaning they are uniquely identified as specific in only a single tissue.

**Figure 2:**
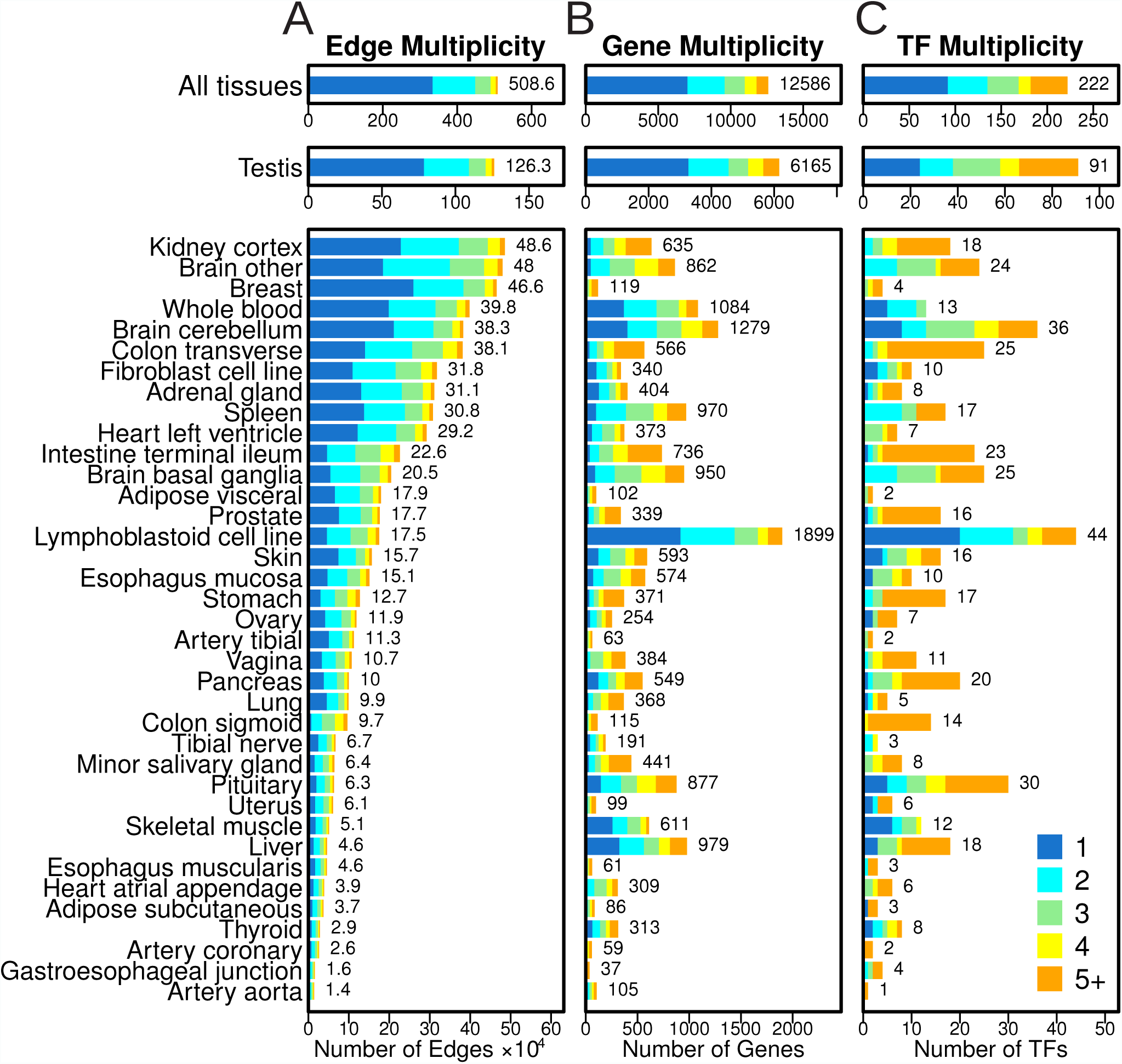
Bar plots illustrating the number of edges (A), genes (B), and transcription factors (TFs, C) that were identified as “specific” to each of the 38 GTEx tissues. The total number of tissue-specific elements identified for each tissue is shown to the right of each bar (edges are shown as a multiple of 10^4^). Tissue-specificity for network elements was defined based on an edge/node having increased weight/expression in one tissue compared to others, thus some edges, genes and TFs were identified as specific to multiple tissues. This “multiplicity” value is indicated by the color of the bars. Note that an edge/gene/TF with a given multiplicity across all tissues (top bar plots) will appear in that number of tissue-specific bar plots (lower bar plots). We found fairly low levels of multiplicity for edges compared to nodes (TFs and genes). TFs also have substantially higher multiplicity compared to genes.

Higher edge multiplicity is often indicative of shared regulatory processes between tissues. For example, 92.2% of sigmoid colon specific edges have a multiplicity greater than one, meaning they are also called specific in other tissues. Further investigation (Supplemental Figure S2A) indicates that 81.0% of these edges are shared with the transverse colon, 46.6% are shared with the small intestine, and 19.3% are shared with the stomach. Similarly, of those edges called specific in the basal ganglia subregion of the brain, 13.3% and 43.3% are also identified as specific in the cerebellum and other subregions of the brain, respectively.

For other tissues the composition of shared edges is quite complex. For example, 78.0% of edges identified as specific in the aorta have a multiplicity greater than one. Of these, the largest fraction is specific to the tibial artery. However, this only includes 14.9% of the aorta-specific edges; additional edges are shared with the testis (12.2%), coronary artery (11.4%), ovary (8.9%), kidney (8.0%), and other brain subregions (7.7%). This shows that even in cases where many of the edges identified as specific in a given tissue have a high multiplicity, as a set, these edges are often distinct from the other tissues.

### 2.2. Identifying Tissue-Specific Network Nodes

Since most analyses of tissue-specificity have examined gene expression, we wanted to know whether the patterns that we observed for the tissue-specific network edges could also be found in tissue-specific expression information. We identified tissue-specific network nodes (TFs and their target genes) using a process analogous to the one we used to identify tissue-specific edges. Specifically, we identified a gene (or TF) as tissue-specific if its median expression in a tissue was greater than the median plus two times the interquartile range of its expression across all tissues. This process identified 12, 586 genes as tissue-specific (41.6% of all genes, see Supplemental Figure S1B–C and Supplemental Table 2); 558 of these genes code for transcription factors (30.6% of TFs in our expression data, this includes 222, or 34.5%, of the 644 TFs that we used in constructing our network models, see Supplemental Material and Methods and Supplemental Table 3).

We find that the number of genes and transcription factors identified as tissue-specific based on expression is not correlated with the number of tissue-specific edges (Figure 2B–C, Supplemental Figure S3). We also observe much higher multiplicity levels for network nodes than for the edges (*p <* 10^-15^ for both genes and TFs by two-sample Chi-squared test), indicating that genes and transcription factors are more likely to be identified as “specific” in multiple tissues than are regulatory edges.

As with the edges, node-multiplicity provides insight into shared functions among the tissues. Consistent with previous findings, testis has the largest number of tissue-specific genes [26, 27] and we find that many of the genes identified as specific in other tissues are also identified as specific in the testis (Supplemental Figure S2B). Other shared patterns of expression mirror what we observed among the network edges. For example, genes identified as specific in the basal ganglia brain subregion include those that are also identified as specific in the cerebellum (41.1%), other brain subregions (67.9%), and the pituitary gland (24.6%). Similarly, 50.4% and 31.3% of sigmoid colon specific genes are shared with the transverse colon and the small intestine, respectively. However, these genes also include those identified as specific in the prostate (23.5%), esophagus (22.6% in the muscularis and 14.8% in the gastroesophageal junction), uterus (18.3%), kidney (14.8%), vagina (13.0%), and stomach (13.0%).

The overlap of genes identified as specific in multiple tissues is quite complex and there are many cases of shared expression patterns between tissues that are not reflected in the tissue-specific network edges we had previously identified. This is especially true for the transcription factor regulators in our network model (Supplemental Figure S2C). For example, only a single transcription factor (*TBX20*) was identified as tissue-specific in the aorta based on our expression analysis. This transcription factor [28, 29] has a high level of multiplicity and was also identified as specific in the coronary artery, testis, pituitary, and heart (both the atrial appendage and left ventricle regions; see Supplemental Table 3 and Supplemental Figure S2C). We find similar patterns in many of the other tissues, including the coronary artery, subcutaneous adipose, esophagus muscularis, tibial nerve, tibial artery, and the visceral adipose. Each of these tissues has only two or three associated tissuespecific transcription factors and almost all of these transcription factors have a multiplicity greater than one, meaning that they were identified as having relatively higher levels of expression in multiple different tissues.

Directly comparing the number of identified tissue-specific transcription factors and genes reveals that there are significantly fewer tissue-specific transcription factors than one would expect by chance (*p* = 2.0 *·* 10^-4^ by two-sample Chi-square test). In addition, transcription factor multiplicity levels are significantly higher than those of genes (*p* = 1.25 *·* 10^-10^ by two-sample Chi-squared test). In other words, TFs are less likely to be identified as tissue-specific compared to genes based on expression profiles. These results imply that tissue-specific regulation may not be due to selective differential expression of transcription factors between tissues.

It should be noted that the transcription factors we identify as tissue-specific based on the GTEx expression data are substantially different than those listed in a previous publication [2] (see Supplemental Figure S4A–C) and used in other GTEx network evaluations [11]. In direct contrast to the results from this previous publication, we find that transcription factors are expressed at higher levels than non-TFs (compare Figure 3A in [2] to Supplemental Figure S4D). This is likely due to technical differences in measuring the expression levels of genes between the two studies. Although state-of-the-art at the time, the data used in the previous publication contained only two samples per tissue and was based on a microarray platform that only assayed expression for a subset of the genes used in our analysis (Supplemental Figure S4E). The differences we find with this previous work highlight the importance of the GTEx project and the opportunity it gives us to revisit our understanding of the role of transcription factors in mediating tissue-specificity.

**Figure 3:**
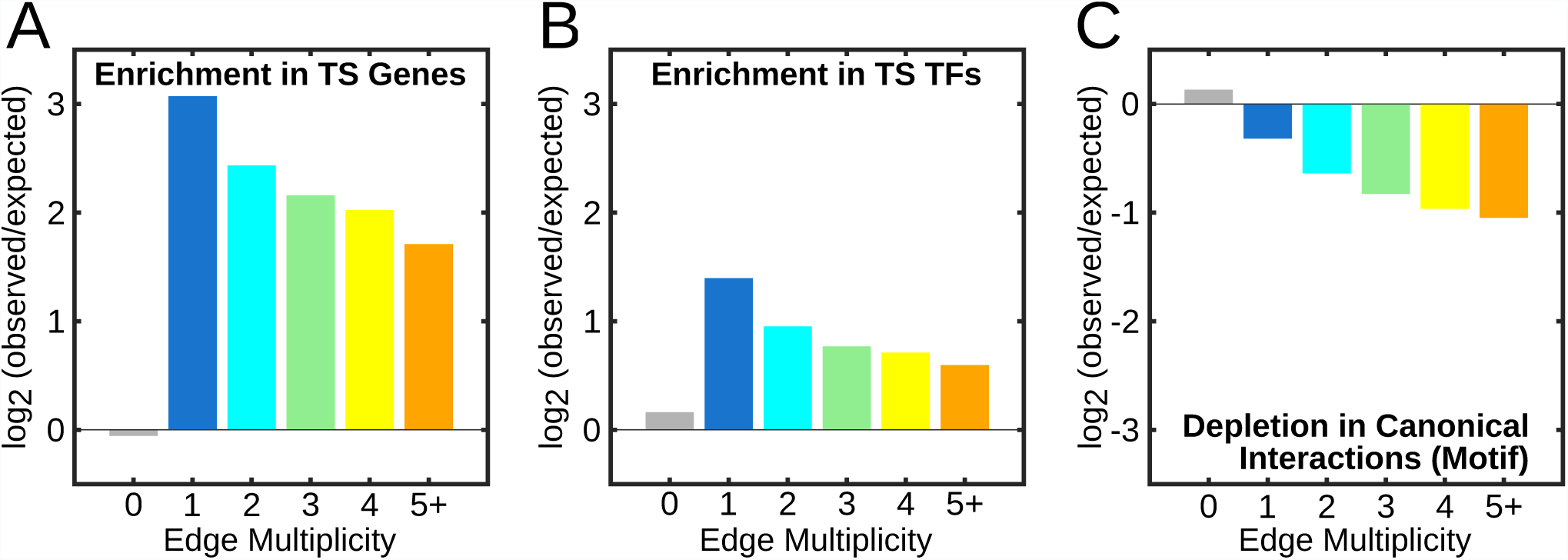
Enrichment of tissue-specific edges in (A) tissue-specific target genes, (B) tissue-specific transcription factors, and (C) canonical transcription factor interactions. For tissue-specific edges of different multiplicities (0 = non-tissue-specific), the log2 of the number of observed/expected number of connections is given.

### 2.3. Characterizing Relationships between Tissue-Specific Network Elements

We tend to think about tissue-specificity in terms of gene expression. However, we know that gene expression arises from a complex set of regulatory interactions between transcription factors and their target genes. The networks inferred from the GTEx data provide us with a unique opportunity to characterize the relationship between the tissue-specific elements—edges, genes, and transcription factors—that help to define tissue phenotype and function.

To do this, we first determined the number of tissue-specific nodes (genes and transcription factors) that are connected to at least one tissue-specific edge. Overall, we found approximately 56% of tissue-specific genes are directly connected to at least one tissue-specific edge (Supplemental Table 4), meaning that tissue-specificity in gene expression is generally associated with tissue-specific changes in regulatory processes. In contrast, tissue-specific transcription factors are *always* connected to at least one tissue-specific edge, meaning that they are always associated with a tissue-specific regulatory process. In fact, we found that nearly every transcription factor is associated with at least one tissue-specific edge in all 38 tissues. This suggests that even transcription factors that are similarly expressed across tissues, and thus not identified as tissue-specific, may play an important role in mediating tissue-specific regulation.

We next quantified the association of tissue-specific edges with tissue-specific nodes. We did this by counting the number of tissue-specific edges that target a tissue-specific gene, summing over all 38 tissues, and dividing by the number one would expect by chance (Supplemental Materials and Methods). We found very high enrichment for tissue-specific edges targeting tissue-specific genes, especially for the most specific edges those with lower multiplicity values (Figure 3A). We repeated this calculation to evaluate whether tissue-specific edges tended to originate from tissue-specific transcription factors. Although we again observed strong enrichment (Figure 3B), this was substantially lower than the enrichment we observed between tissue-specific edges and genes.

Finally, because PANDA leverages multiple sources of data to reconstruct its models, we analyzed tissue-specific edges in the context of both the input co-expression data and the canonical transcription factor-target gene interactions (defined by the presence of a TF motif in the promoter region of a target gene) we used to seed our networks. We found that tissue-specific edges are distinct from those identified using only co-expression information (Supplemental Figure S2D). In addition, tissue-specific edges are depleted for canonical transcription factor interactions (Figure 3C). Thus, many of the tissue-specific regulatory interactions we identified using PANDA would have been missed if we had relied solely upon co-expression or transcription factor motif targeting information to define a regulatory network.

### 2.4 Evaluating Tissue-Specific Regulation of Biological Processes

As noted previously, we find that transcription factors are less likely than other genes to be identified as tissue-specific based on their expression profile, and even those identified as tissue-specific tend to have a high multiplicity (are specific in multiple tissues). In addition, although tissue-specific transcription factors are significantly associated with tissue-specific network edges, this association is much lower than the one between tissue-specific genes and edges. These results led us to hypothesize that both tissue-specific and non-tissue-specific transcription factors (as defined based on expression information) play an important role in mediating tissue-specific biological processes.

We selected one of the brain tissue subregions (“Brain other”) to test this hypothesis since this tissue had one of the largest number of tissue-specific edges (after testis and kidney) and the majority of genes and transcription factors called as specific to this tissue are also specific in other tissues (have high multiplicity, see Figure 2). We ran a pre-Ranked Gene Set Enrichment Analysis (GSEA) [30] on each transcription factor’s tissue-specific targeting profile to evaluate the role of transcription factors in regulating particular biological processes (see Supplemental Material and Methods).

Figure 4A shows the Gene Ontology (GO) Biological Process terms that were significantly enriched (*F DR <* 0.001; GSEA Enrichment Score, *ES >* 0.65) for tissue-specific targeting by at least one transcription factor in this brain tissue subregion. Among the significant processes are many brain-related functions, including axonogenesis, axon guidance, regulation of neurogenesis, regulation of neurotransmitter levels, and neurotransmitter secretion. A hierarchical clustering (Euclidean distance, complete linkage) of the GSEA enrichment profiles indicates that transcription factor regulators are generally associated with either an increased or decreased targeting of genes involved in these brain-associated processes. To our surprise, the transcription factors that are positively associated with brain-related functions are not any more likely to be expressed in a tissue-specific manner than transcription factors that are not positively associated with these functions.

**Figure 4:**
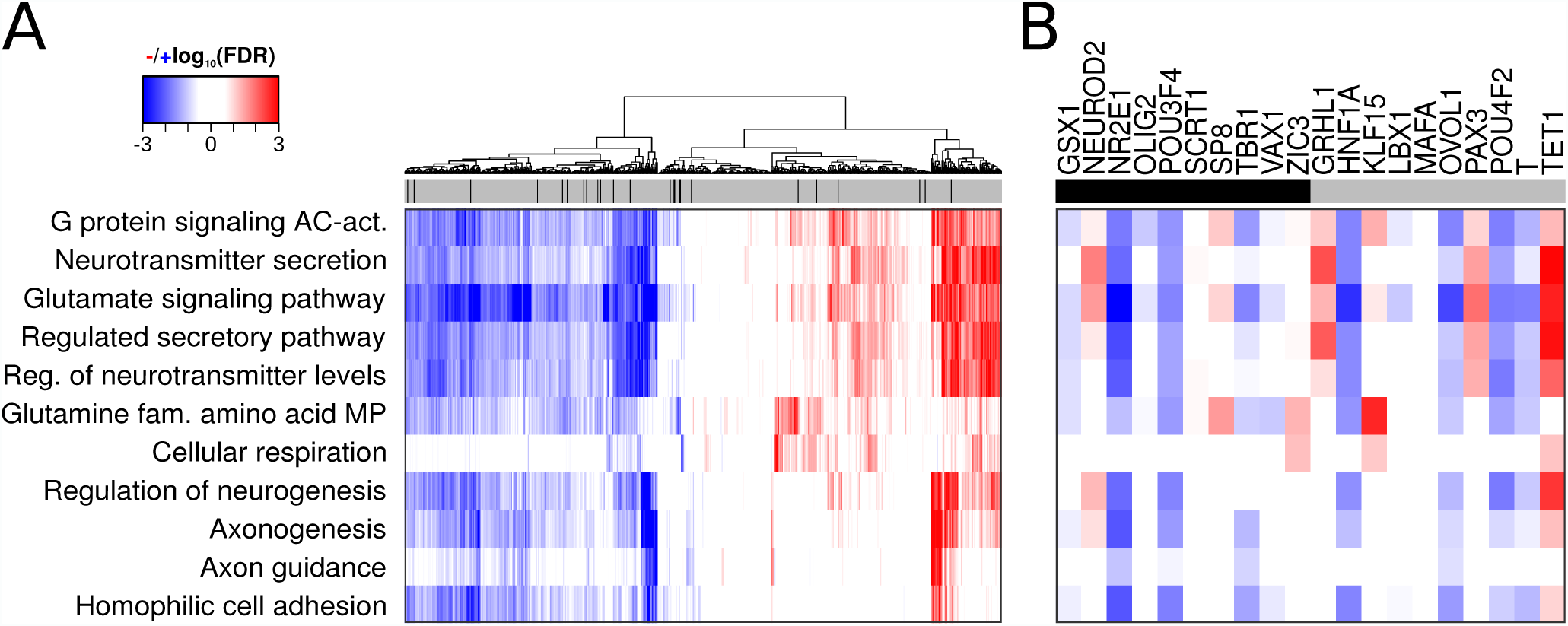
(A) Heatmap depicting the GSEA results for the targets of 644 transcription factors in the “Brain other” gene regulatory network model. The figure includes all significantly targeted (*F DR <* 0.001, GSEA Enrichment Score, *ES >* 0.65) GO Terms. Positive enrichment scores, indicating increased targeting of genes by a given transcription factor, are shown in red. Negative scores are in blue. FDR values greater than 0.25 appear white. The top bar indicates whether a transcription factor was also identified as specific (black) to “Brain other” or not (gray). (B) Heatmap for the ten most (black) and ten least (gray) tissue-specific transcription factors. AC: adenylate cyclase, act.: activating pathway, reg. : regulation, fam.: family, MP: metabolic process.

To ensure this result was not due to the threshold we used when identifying tissue-specific TFs, we selected the ten transcription factors with the highest and lowest expression enrichment in this brain-tissue subregion (see Supplemental Material and Methods) and performed a detailed investigation of their GSEA profiles (Figure 4B). NEUROD2 and SP8 were the top tissue-specific transcription factors with brain-function associated targeting profiles; these TFs play important roles in brain function [31–33]. In addition, four of the highly non-tissue-specific transcription factors (based on expression)—GRHL1, KLF15, PAX3, and TET1—have positive enrichment for targeting genes with relevant brain functions. These non-brain-specific transcription factors have been shown to play an important role in neuroblastoma [34], neuronal differentiation [35], brain development [36], and neuronal cell death [37], respectively.

Finally, we identified 38 transcription factors that exhibit highly significant (*F DR <* 0.001 and *ES >* 0.65) differential targeting of the identified functions. Only one of these transcription factors (RFX4) was also identified as tissue-specific based on expression analysis. When we repeated this analysis for all 38 tissues we found similar patterns, with low overlap between the transcription factors identified as tissue-specific based on expression and those that have strong patterns of differential targeting (Supplemental Figure S5 and Supplemental Table 5). These results indicate that transcription factors do not have to be differentially expressed to play significant tissue-specific regulatory roles. Rather, changes in their targeting patterns allow them to regulate tissue-specific biological processes.

### 2.5 Tissue-Specific Organization of Biological Processes

Because of the high level of multiplicity that we previously observed, especially for transcription factors (see Figure 2), we next examined shared functional regulation based on tissue-specific targeting patterns. Specifically, we ran GSEA on the tissue-specific targeting profile of each transcription factor in each of the 38 tissues and selected GSEA results that represented highly significant positive enrichment for tissue-specific TF-targeting (*F DR* < 0.001 and *ES* > 0.65; all results contained in Supplemental Table 6). We then clustered these associations [38] (see Supplemental Material and Methods) and identified 48 separate “communities,” or groups of GO terms associated with TF/tissue pairs [39, 40] (Figure 5A). Properties of the identified communities, including the number of terms, TFs, and tissues represented in each, are included in Supplemental Table 7.

**Figure 5:**
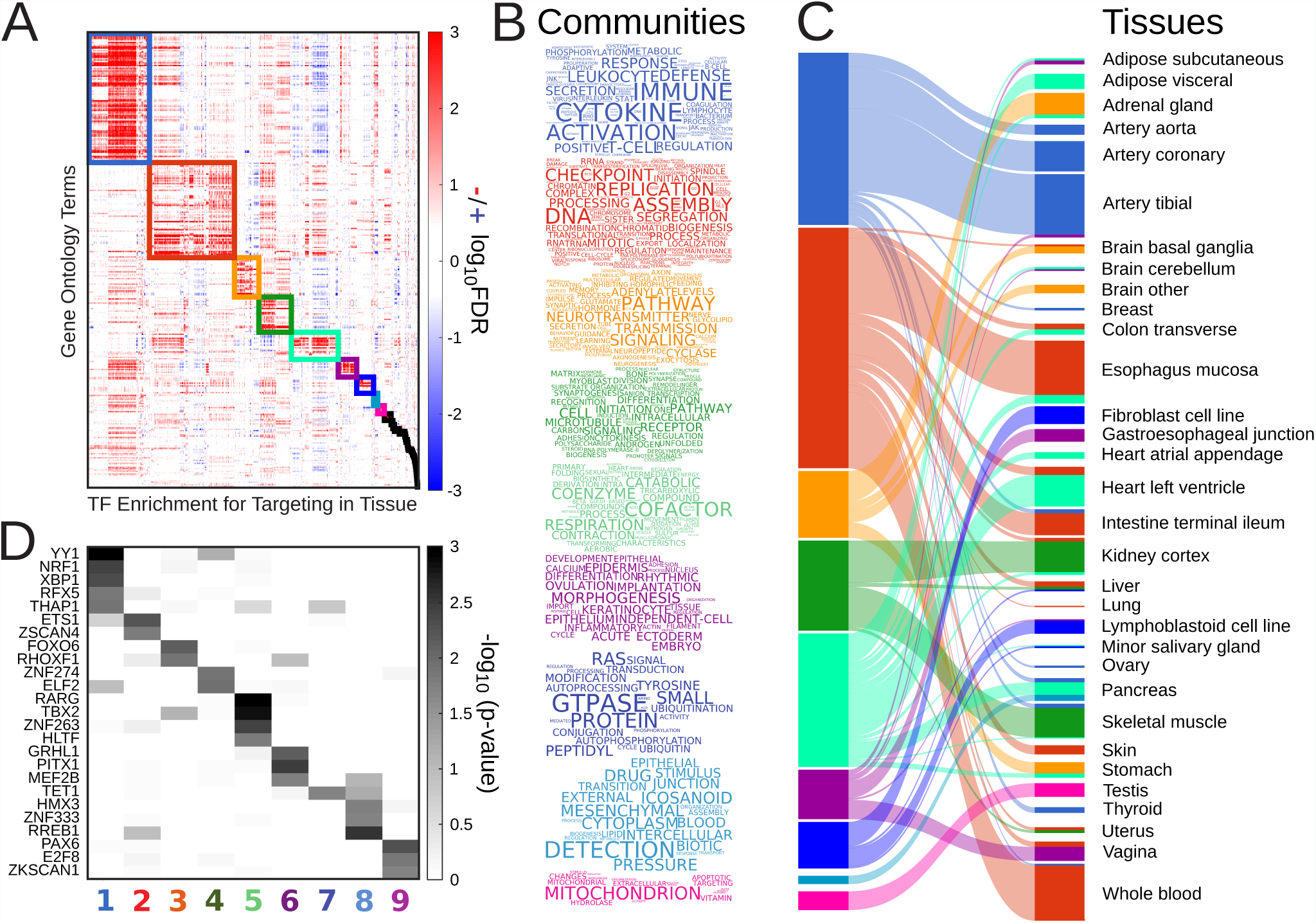
(A) Heatmap depicting communities of GO terms that were significantly targeted (*F DR <* 0.001, GSEA Enrichment Score *ES >* 0.65) based on a GSEA analysis run on all possible tissue-transcription factor pairs. Tissue-transcription factor pairs were also clustered and identified with each community. (B) Word clouds summarizing the processes contained in each community. (C) An illustration of the tissues associated with each community. Edge width indicates the number of transcription factors that were identified as differentially targeting at least one signature in the community in a particular tissue. For simplicity we only illustrate the top nine communities (left) and connections to tissues that include five or more transcription factors. For an interactive version of figure C, see Supplemental File 1. (D) Heatmap of the top transcription factors involved in targeting the nine largest communities; the grayscale gradient represents the probability that a transcription factor would be associated with a community by chance given a random shuffling of community assignments (see Supplemental Materials and Methods).

Nine communities had more than five associated GO terms. Further inspection showed that these communities often included sets of highly related functions, such as those associated with immune response (Community 1), cell proliferation (Community 2), synaptic transmission (Community 3), extracellular structure (Community 4), cellular respiration (Community 5), ectoderm development (Community 6), protein modification (Community 7), cellular response (Community 8), and the mitochondrion (Community 9).

We used word clouds to summarize this information and provide a snapshot of the functions associated with each of these nine communities (Figure 5B; Supplemental Materials and Methods). We also examined what tissues were associated with each community and found that communities were generally dominated by enrichment for increased functional targeting in a select set of tissues (Figure 5C). For example, Community 1 is highly associated with the tibial and coronary arteries, Community 3 is highly associated with two of the brain subregions (“Brain other” and “Brain basal ganglia”) as well as the adrenal gland and stomach, and Community 4 is highly associated with skeletal muscle, as well as the kidney cortex. Although some of the communities represent sets of functions that are common to multiple tissues, these associations make biological sense. For example, some tissues, such as skin and whole blood, have higher rates of proliferation compared to others and so we might expect increased targeting of cell cycle functions in these tissues.

The remaining 39 communities had five or fewer GO term members but often capture important associations between tissues and biological function (Supplemental Figure S6). For example, Community 17 contains two GO Biological Processes term members, “spermatid development” and “spermatid differentiation” and is enriched for positive tissue-specific targeting in the testis (12 TFs). Community 21 contains exactly one term member, “digestion” and is enriched for positive tissue-specific targeting in the sigmoid colon (25 TFs). Community 25 also contains exactly one GO term, “hormone metabolic process” and is enriched for positive tissue-specific targeting in the pituitary (7 TFs, including FEZF1, HOXA13, NRL, POU3F4, SIX3, SOX2, and SRY). The GO term and TF/tissue members of all communities are contained in Supplemental Table 7.

In addition to identifying tissue-specific function, we identified several transcription factors that appear to mediate similar biological functions across multiple tissues (Figure 5D; Supplemental Materials and Methods). For example, Community 1 (immune response) includes targeting profiles from 340 different TFs and 23 tissues (Supplemental Table 7). Further inspection reveals that five of these transcription factors have significantly more associations in Community 1 than one would expect by chance. These transcription factors include RFX5, which plays an essential role in the regulation of the major histocompatibility complex class II (MHC-II), a key component of the adaptive immune system [41], and YY1, which was recently reported to inhibit differentiation and function of regulatory T cells [42].

### 2.6. Maintenance of Tissue-Specificity in the Global Regulatory Framework

The analysis we have presented thus far has focused primarily on tissue-specific network edges, or regulatory interactions that have an increased likelihood in one, or a small number of tissues, compared to others. However, we know that these tissue-specific interactions work within the context of a larger “global” gene regulatory network, much of which is the same in many tissues. Therefore, we investigated how tissue-specific regulatory processes are reflected in changes to the overall structure and organization of each tissue’s “global” gene regulatory network.

To begin, we analyzed the connectivity of nodes separately in each of the 38 tissues’ gene regulatory networks using two measures: (1) degree, or the number of edges connected to a node, and (2) betweenness [43], or the number of shortest paths passing through a node (Figure 6A). Although both of these measures are well established in the field of network science, betweenness in particular has only occasionally been used to analyze biological networks [11, 44], and, to our knowledge, has not been used to assess tissue-specific gene regulatory networks. Because of the complete nature of the networks estimated by PANDA, we used algorithms that account for edge weight when calculating these measures [45] (see Supplemental Materials and Methods). For each tissue, we then compared the median degree and betweenness values of tissue-specific genes to the median degree and betweenness values of non-tissue-specific genes (Figure 6B).

**Figure 6:**
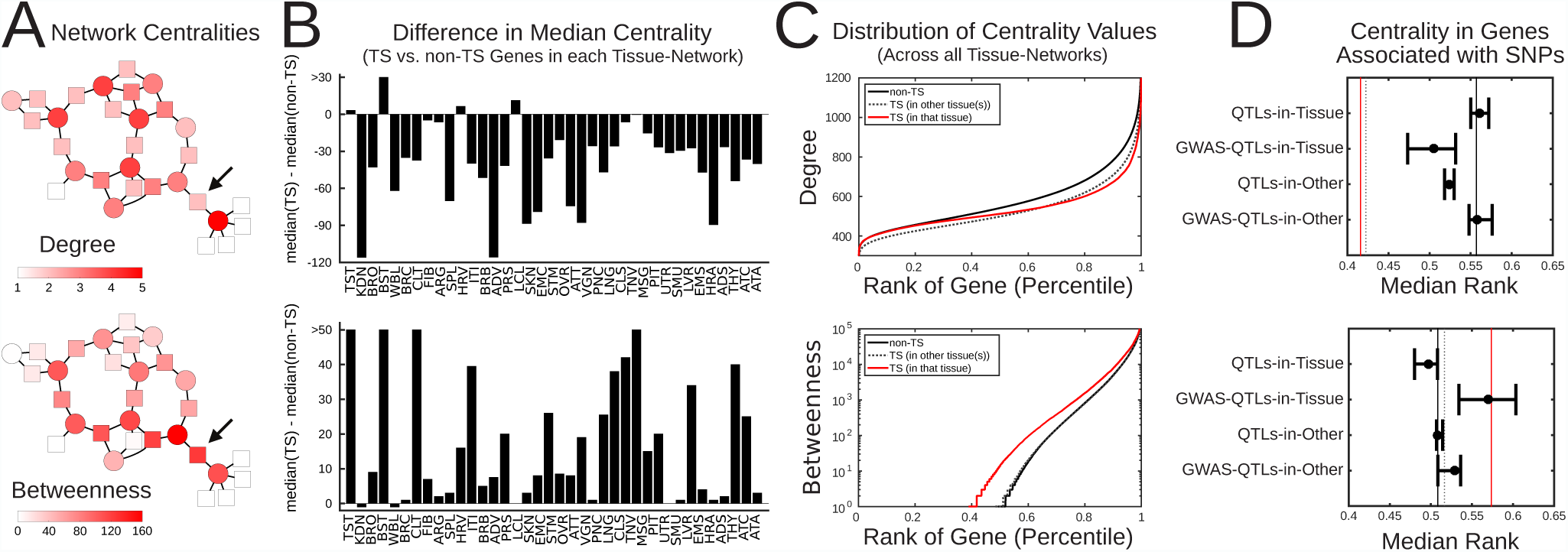
(A) An example network illustrating the difference between high degree and betweenness. Transcription factors are shown as circles and target genes as squares. The color of each node indicates its centrality based on the relevant measure. An example node is shown with low degree but high betweenness. (B) Difference in the median centrality of tissue-specific genes compared to non-tissue-specific genes in each of the 38 networks. (C) Distribution of centrality values for all non-tissue-specific genes (black), genes specific in a particular tissue (red), and genes called tissue-specific in some tissue, but not the tissue of interest (gray dashed line). (D) The median and interquartile range across tissues of the percentile-rank of genes (based on their centrality, as in panel C) that have an eQTL association in that tissue (QTLs-in-tissue), have an eQTL association with a GWAS variant in that tissue (GWAS-QTLs-in-Tissue), have an eQTL association in another tissue (QTLs-in-other) or have an eQTL association with a GWAS variant in another tissue (GWAS-QTLs-in-Other). For comparison, the median rank of tissue-specific and non-tissue-specific genes across these tissues is indicated by a red and black line, respectively. Note that the analysis shown in this panel is limited to 19 tissues and the 29, 155 genes considered in both this analysis and the eQTL analysis described in [46].

This analysis showed that tissue-specific genes generally have a lower degree than non-tissue-specific genes. This may initially seem contradictory to our observation that tissue-specific genes are highly targeted by tissuespecific edges (Figure 3A). However, we also found that tissue-specific edges tended to be associated with noncanonical regulatory events (Figure 3C), which generally have lower weights in our network models. The analysis presented here considers all regulatory interactions (both tissue-specific and non-tissue-specific) leading to a network whose structure is largely dominated by canonical regulatory events. Thus, we can conclude that tissuespecific genes gain targeting from tissue-specific edges, consistent with our previous finding. However, in the context of the global gene regulatory network, the targeting of these tissue-specific genes is much lower as compared to other, non-tissue-specific genes [47].

These findings are consistent with the notion that processes required for a large number of (or all) tissues need to be stably regulated. Thus one might expect these to be more tightly controlled and therefore central to the network. Indeed, when we examine the distributions of degree values (Figure 6C) we find the largest differences are between tissue-specific and non-tissue-specific genes with high degree (network hubs), with a bias for non-tissue-specific genes to have high degree values. In other words, we observe a depletion of tissue-specific genes among the gene regulatory network hubs.

Our analysis also showed that tissue-specific genes have higher median betweenness compared to non-tissue-specific genes. This indicates that tissue-specific func tion may be mediated by tissue-specific regulatory paths through the global network structure, creating new avenues by which information can “flow” through tissuespecific genes (as measured by betweenness) despite their relatively low overall connectivity (as measured by degree). Indeed, when we examine the distribution of betweenness values (Figure 6C), we find that tissue-specific genes are significantly enriched for small but measurable values, while non-tissue-specific genes are more likely to have no shortest paths running through them (*p* < 10^-15^ by one-sided Kolmogorov-Smirnov test). We note that the signals we observe here are absent in a network constructed solely based on canonical transcription factor-target gene interactions (Supplemental Figure S7) suggesting that these regulatory paths are most likely the result of the inclusion of tissue-specific edges in the global regulatory network structure.

### 2.7. Implications of Tissue-Specific Regulation

One important reason for modeling tissue-specific regulatory networks is to provide a baseline that can be used to better understand how regulatory processes might be perturbed by disease, or in the presence of other biological factors, such as a genetic variant. To evaluate the utility of our tissue-specific networks in the context of this type of information, we leveraged information regarding tissue-specific, *cis*-acting expression quantitative trait loci (eQTLs) that we had previously identified using the GTEx data [46], together with information from Genome Wide Association Studies (GWASes), as curated in the GWAS catalog (http://www.ebi.ac.uk/gwas/). For this analysis, we focused on 19 tissues for which we had reliable estimates of *cis*-eQTLs, and the 29, 155 genes that were considered when performing the eQTL analysis and that were included in our network models. For more information see Supplemental Materials and Methods.

To begin, for each tissue, we identified which genes had at least one significant (*F DR* < 0.05) eQTL association. 25, 819 genes (88.6%) had at least one eQTL association in at least one of the tissues (from 3, 317 in brain-other to 10, 997 in thyroid). As a group, these QTL-associated genes were significantly enriched for transcription factors (*p* = 9.09 *·* 10^-5^ by hypergeometric probability), but significantly depleted for tissue-specific genes (*p* < 10^-15^ by hypergeometric probability). However, when we evaluated whether genes that are specific to a particular tissue are also more likely to have an eQTL association in that same tissue, we observed a significant enrichment (*p* = 1.69 *·* 10^-14^ by hypergeometric probability). This indicates that, although tissue-specific genes as a *group* are less likely to have associations with genetic variants, when they do, it is within the context of their tissue environment.

Next, to understand if these findings might have disease relevance, we focused on the subset of genes that had a significant *cis*-eQTL association with one of the genetic variants listed in the GWAS catalog. Only 308 genes (1%) had one or more eQTL associations with a GWAS-variant (Supplemental Table 8). Of these 308 genes, only 39 (12.7%) were also identified as tissue-specific (significantly depleted; *p* = 1.51 · 10^-4^ by hypergeometric probability). In contrast to the QTL-associated genes, the GWAS-associated subset were neither enriched nor depleted for transcription factors. In addition, when we investigated whether the eQTL associations for these genes tended to occur in the same tissue in which those genes were identified as specific, we observed significant depletion (*p* = 5.46 *·* 10^-4^ by hypergeometric probability).

Finally, to understand how these findings might be reflected in the context of our gene regulatory networks, we evaluated the centralities of these genes, both within the tissue for which they had the identified association(s), and across all other tissues (those in which they did not have any significant eQTL associations). Specifically, we ranked genes in each tissue based on their centrality, found the median rank of the QTL-associated and GWAS-QTL-associated genes in both the tissue where they had a significant association, and in each of the other tissues. We plot the range of these median values across the 19 examined tissues in Figure 6D.

We see a clear signal for variant-associated genes in both their degree and betweenness. In particular, genes that are associated with a GWAS variant have lower degree and higher betweenness in their corresponding tissue network as compared to the set of genes that have any eQTL association in that tissue. Furthermore, this behavior is distinct in their corresponding tissue-network as opposed to other tissue-networks. We note that this increase in betweenness coupled with a decrease in degree is exactly the same pattern as we observed for tissue-specific genes. However, interestingly, as we noted above, GWAS-QTL-associated genes are highly depleted for tissue-specificity. This may help explain why many GWAS loci are associated with multiple diseases. It also suggests that our network models are capturing important routes of regulatory information flow beyond those necessary for the maintenance of tissue-specific processes and thus have the potential to be used to understand disease-related regulatory information.

## 3. DISCUSSION

We used gene expression data from GTEx, together with other sources of regulatory information, to reconstruct and characterize regulatory networks for 38 tissues and to assess tissue-specific gene regulation. We used these networks to identify tissue-specific edges and used the gene expression data to identify tissue-specific nodes (transcription factors and genes). We found that, although tissue-specific edges are enriched for connections to tissue-specific transcription factors and genes, they are also depleted for “canonical” interactions (defined based on a transcription factor binding site in the target gene’s promoter). In addition, edges are often uniquely called as specific in only one tissue while tissue-specific genes often have a high “multiplicity,” meaning that they were identified as specific in more than one tissue.

In particular, we found that genes that encode for transcription factors were especially likely to be identified as specific in multiple different tissues. This suggests that the notion of a “tissue-specific” transcription factor based on expression information should be considered with care, especially in the context of transcriptional regulation. Indeed, analysis of tissue-specific targeting patterns in our regulatory networks indicated that transcription factor expression is not the primary driver of tissue-specific functions. Our network analysis found many transcription factors that are known to be involved in important tissue-specific biological processes that were not identified as tissue-specific based on their expression profiles. These findings are consistent with what we might expect [48]. There are approximately 30, 000 genes in the human genome, but fewer than 2, 000 of these encode transcription factors [2] (of which we analyzed only 644—those with high quality motif information). Given the large number of tissue-specific functions that must be regulated, it makes sense that changes in complex regulatory patterns are responsible for tissue-specific gene expression, not the activation or deactivation of individual regulators.

Our results suggest that transcription factors primarily participate in tissue-specific regulatory processes via alterations in their targeting patterns. To understand the regulatory context of these tissue-specific alterations, we investigated the topology of each of the 38 global tissue regulatory networks (containing information for all possible edges). We found that tissue-specific genes generally are less targeted (have a lower degree) than nontissue-specific genes. However, tissue-specific genes exhibit an increase in the number of regulatory paths running through them (have a higher betweenness) as compared to non-tissue-specific genes. These results suggest that tissue-specific regulation does not occur in dense portions of the regulatory network, or by the formation of tissue-specific hubs. Rather, tissue-specific genes are central to the regulatory network on an intermediate scale due to the influence of tissue-specific tissue-specific regulatory paths [47]. We believe this result supports the notion that tissue-specific function is largely driven by non-canonical interactions. Such interactions could, for example, be interactions through TF complexes (no direct binding between a TF to the promoter of its target gene), binding of a TF to an alternative motif, or interactions outside of a gene’s promoter (for example binding to an enhancer) [49].

Overall, our analysis provides a more comprehensive picture of tissue-specific regulatory processes than reported previously. Our comparison of global gene regulatory network models across a large set of human tissues provided important insights into the complex regulatory connections between genes and transcription factors, allowed us to identify how those structures are subtly different in each tissue, and ultimately led us to better understand how transcription factors regulate tissue-specific biological processes. One important result from our analysis is that transcription factor expression information is very poorly correlated with tissue-specific regulation of key biological functions. At the same time, we find that alterations in transcription factor targeting cause a shift in the structure of each tissue’s regulatory network, such that tissue-specific genes occupy central positions by virtue of tissue-specific regulatory paths that run through the global network structure.

Taken together, these results support the notion that tissue-specificity likely arises from adjusting and adapting existing biological processes. In other words, tissue-specific biological function occurs as a result of building on an existing regulatory structure such that both common and tissue-specific processes share the same underlying network core. Ultimately, our work suggests that regulatory processes need to be analyzed in the context of specific tissues, particularly if we hope to understand disease and development, to develop more effective drug therapies, and to understand the potential side effects of drugs outside of the target tissue. It also establishes a framework in which to think about the evolution of tissue-specific functions, one in which new processes are integrated into an established gene regulatory framework.

## 4. ACKNOWLEDGMENTS

This work was supported by grants from the US National institutes of Health, including grants from the National Heart, Lung, and Blood Institute (5P01HL105339, 5R01HL111759, 5P01HL114501, K25HL133599), the National Cancer Institute (5P50CA127003, 1R35CA197449, 1U01CA190234, 5P30CA006516, P50CA165962), the National Institute of Allergy and Infectious Disease (5R01AI099204), and the Charles A. King Trust Post-doctoral Research Fellowship Program, Sara Elizabeth O’Brien Trust, Bank of America, N.A., Co-Trustees. Additional funding was provided through a grant from the NVIDIA foundation. This work was conducted under dbGaP approved protocol #9112 (accession phs000424.v6.p1).

## 5. AUTHOR CONTRIBUTIONS

All authors conceived of the study; ARS, JP, MF, JNP, KG, and MLK analyzed the data; ARS, MLK, and KG drafted the initial manuscript. All authors contributed to the reviewing and editing of the manuscript. All authors read and approved the final manuscript.

## SUPPLEMENTAL MATERIALS AND METHODS

### S.1. GTEx RNA-Seq Data

We downloaded the Genotype-Tissue Expression (GTEx) version 6.0 RNA-Seq data set (phs000424.v6.p1, 2015-10-05 released) from dbGaP (approved protocol #9112). GTEx release version 6.0 sampled 551 donors with phenotypic information and included 9, 590 RNA-Seq assays. GTEx assayed expression in 30 tissue types, which were further divided into 53 tissue subregions (51 tissues and two derived cell lines) [1]. After removing tissues with very few samples (fewer than 15), we were left with 27 tissue types from 49 subregions. Using YARN (http://bioconductor.org/packages/release/bioc/html/yarn.html) we performed quality control, gene filtering, and normalization preprocessing. Briefly, we performed principal coordinate analysis (PCoA) using Y-chromosome genes to test for sample sex misidentification; we identified and removed GTEX-11ILO which was annotated as female but clustered with the males and was later confirmed to be an individual who underwent sex reassignment surgery (Kristin Ardlie, Broad Institute, private communication). We also used principal coordinate analysis on autosomal genes to group related body regions that had highly sim ilar gene expression profiles. For example, skin samples from the lower leg (sun exposed) and from the suprapubic region (sun unexposed) shared gene expression profiles and were grouped as “skin,” while the transverse and descending colon were very different and were retained as distinct tissues. Gene expression data were then normalized using qsmooth [2] which performs a sparsity aware normalization that provides comparable expression profiles across all tissues. This preprocessing resulted in a dataset of 9, 435 gene expression profiles assaying 30, 333 genes in 38 tissues from 549 individuals. More detailed information on the normalization process and a complete description of the 38 final tissues and the associated samples are described elsewhere [3]. Consistent with GTEx, genes are denoted by their Ensembl IDs.

### S.2. Regulatory Network Reconstruction

We used the PANDA (Passing Attributes between Networks for Data Assimilation) network reconstruction algorithm [4] to estimate gene regulatory networks in each of the 38 GTEx tissues (see Section S.1). PANDA incorporates regulatory information from three types of data: gene expression (used to create a co-expression network), protein-protein interaction, and a “prior” network based on mapping transcription factors to their putative target genes (used to initialize the algorithm).

#### Additional Gene Expression Data Processing

We filtered the normalized GTEx gene expression data (see above) to retain only the 30, 243 genes that also had a significant motif-hit in their promoter region (see below). These genes were used when constructing our regulatory network models.

#### Prior Regulatory Network Based on Transcription Factor Motif Information

To create a “prior” regulatory network between transcription factors and genes, we downloaded Homo sapiens transcription factor motifs with direct/inferred evidence from the Catalog of Inferred Sequence Binding Preferences CIS-BP (http://cisbp.ccbr.utoronto.ca, accessed: July 7, 2015). For each unique transcription factor, we selected the motif with the highest information content, resulting in a set of 695 motifs. We mapped these transcription factor position weight matrices (PWM) to the human genome (hg19) using FIMO [5] and retained highly significant matches (*p* < 10^-5^) that occurred within the promoter regions of Ensembl genes (GRCh37.p13; annotations downloaded from http://genome.ucsc.edu/cgi-bin/hgTables, accessed: September 3, 2015); promoter regions were defined as [–750, +250] around the transcription start site (TSS). After intersection to only include genes and transcription factors with expression data (see above) and at least one significant promoter hit, this process resulted in an initial map of potential regulatory interactions involving 644 transcription factors targeting 30, 243 genes.

#### Prior Protein-Protein Interaction Network

We estimated an initial protein-protein interaction (PPI) network between all transcription factors (TFs) in our motif prior using interaction scores from StringDb v10 (http://string-db.org, accessed: October 27, 2015). PPI interaction scores were divided by 1, 000 and selfinteractions were set equal to one.

#### Recontructing Networks using PANDA

For each of the 38 tissues, we used the GTEx gene expression data to calculate pairwise co-expression levels (based on Pearson correlation) between the 30, 243 target genes. We then used PANDA to combine this information with the prior regulatory network and protein-protein interaction network. This produced 38 regulatory networks, one for each tissue, with edges predicted between 644 transcription factors and 30, 243 target genes. PANDA returns complete, bipartite networks with edge weights similar to z-scores that represent the likelihood of a regulatory interaction. We transformed these z-scores to positive values using:

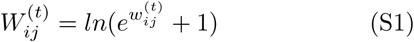

where 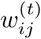 is the edge weight calculated by PANDA between a TF (*i*) and gene (*j*) in a particular tissue (*t*), and 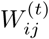 is the transformed edge weight. These transformed edge weights are positive and so avoid issues related to calculating centrality measures on graph with negative edge weights (see Section S.10).

### S.3. Quantification of Tissue-Specificity vs Generality of Network Edges

Each of the 38 reconstructed PANDA networks contains scores, or “edge weights,” for every possible transcription factor-to-gene interaction (see Section S.2). We used these edge weights to identify tissue-specific network edges. To do this, we compared the weight of an edge between a transcription factor (*i*) and a gene (*j*) in a particular tissue (*t*) to the median and interquartile range (IQR) of its weight across all 38 tissues:

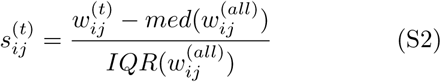

We then defined an edge with an edge specificity score 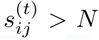 as specific to tissue *t*. We varied the cutoff *N* from 1 to 3, by steps of 0.25. Supplemental Figure S1A shows the fraction of edges that are identified as tissue-specific at each cutoff. We selected a cutoff of *N* = 2 to define tissue-specific edges in order to be consistent with the cutoff used to define tissue-specific nodes (see Section S.4). We also defined the “multiplicity” of an edge as:

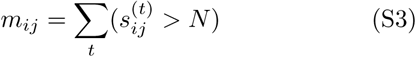

This value represents the number of tissues in which an edge is identified as specific. The overlap in edges identified as specific to each tissue can be found in Supplemental Figure S2A.

### S.4. Quantification of Tissue-Specificity vs Generality of Network Nodes

We wished to know if the tissue-specific edges were a direct reflection of the underlying gene expression data, or if the networks might be providing additional insight into the tissue-specific regulation of genes. Therefore, we identified tissue-specific network nodes (TFs and their target genes) by applying an analogous definition as we used to define tissue-specific edges to the GTEx gene expression data. We compared the median expression level 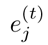 of a gene (*j*) in a particular tissue (*t*), to the median and interquartile range of its expression across all samples:

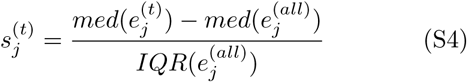

We then defined a gene with gene specificity score 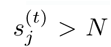 as specific to tissue *t*. We varied the cutoff *N* from 1 to 3, by steps of 0.25. Supplemental Figure S1B shows the fraction of tissue-specific genes identified at each cutoff. Based on this analysis, we selected a cutoff of *N* = 2 because with that cutoff approximately half of all genes are identified as tissue-specific. We also defined the “multiplicity” of a gene as:

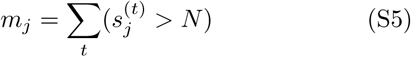

This value represents the number of tissues in which a gene is identified as specific. In Supplemental Figure S1C we show some examples of non-tissue-specific and tissue-specific genes with different levels of multiplicity. We observe that the term “tissue-specific” is largely a misnomer. Many genes have a multiplicity greater than one, meaning that they are not actually “specific” to a particular tissue, but rather have a relatively higher level of expression in a subset of tissues compared to the others. Information regarding the tissue-specificity and multiplicity of genes can be found in Supplemental Table 2. The overlap in genes identified as specific to each tissue can be found in Supplemental Figure S2B.

To identify tissue-specific network nodes, other approaches, such as a differential expression analysis using Limma or ANOVA, could have been used. We decided to use the approach described in Equations S4–S5 for two reasons. First, this allowed us to link the tissue-specificity of network nodes to the specificity of network edges (Equations S2–S3). While multiple expression samples were available for each tissue (ranging from 36 to 779 samples, with a median of 210.5 samples), we only had one network available per tissue. We therefore could not use a statistical test that would compare groups of network edges. Second, approaches such as Limma or ANOVA assume that most genes are not differentially expressed between different conditions. In addition, these approaches assume normality. Because the GTEx expression data are not normally distributed, and because we observed global expression differences in expression between the different tissues (with large global expression differences in, for example, testis), these underlying null assumptions are likely not well founded.

**Supplemental Figure S1:**
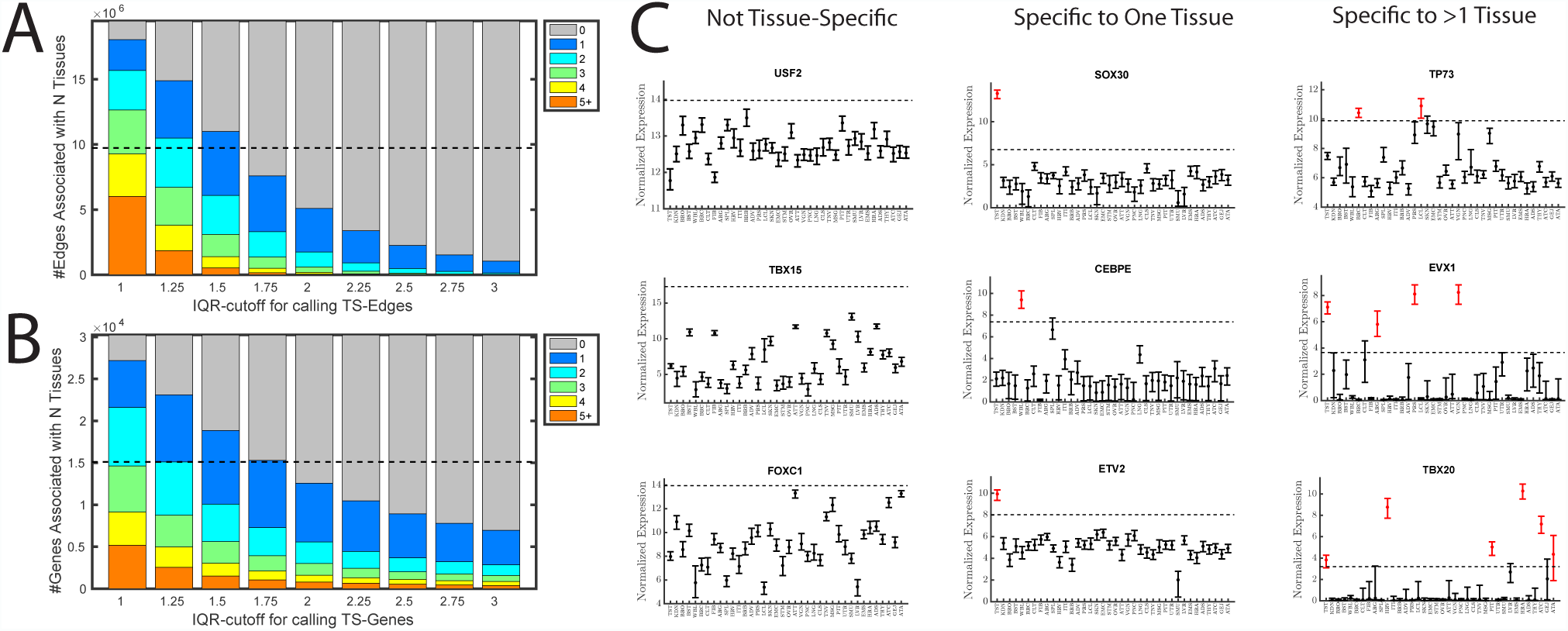
Identification of tissue-specific edges and nodes. (A) Number of edges of a given multiplicity at various cutoffs (*N*). (B) Number of genes of a given multiplicity at various cutoffs (*N*). (C) Examples of genes with various multiplicity levels. For each tissue we show the median (dot) and interquartile range (IQR, range) of a gene’s expression in samples from that tissue. The dashed line indicates the cutoff used to define a gene as tissue-specific (the median plus two times the IQR of expression levels across all samples). A gene is identified as tissue specific if its median expression across samples in a particular tissue is above the cutoff.

**Supplemental Figure S2:**
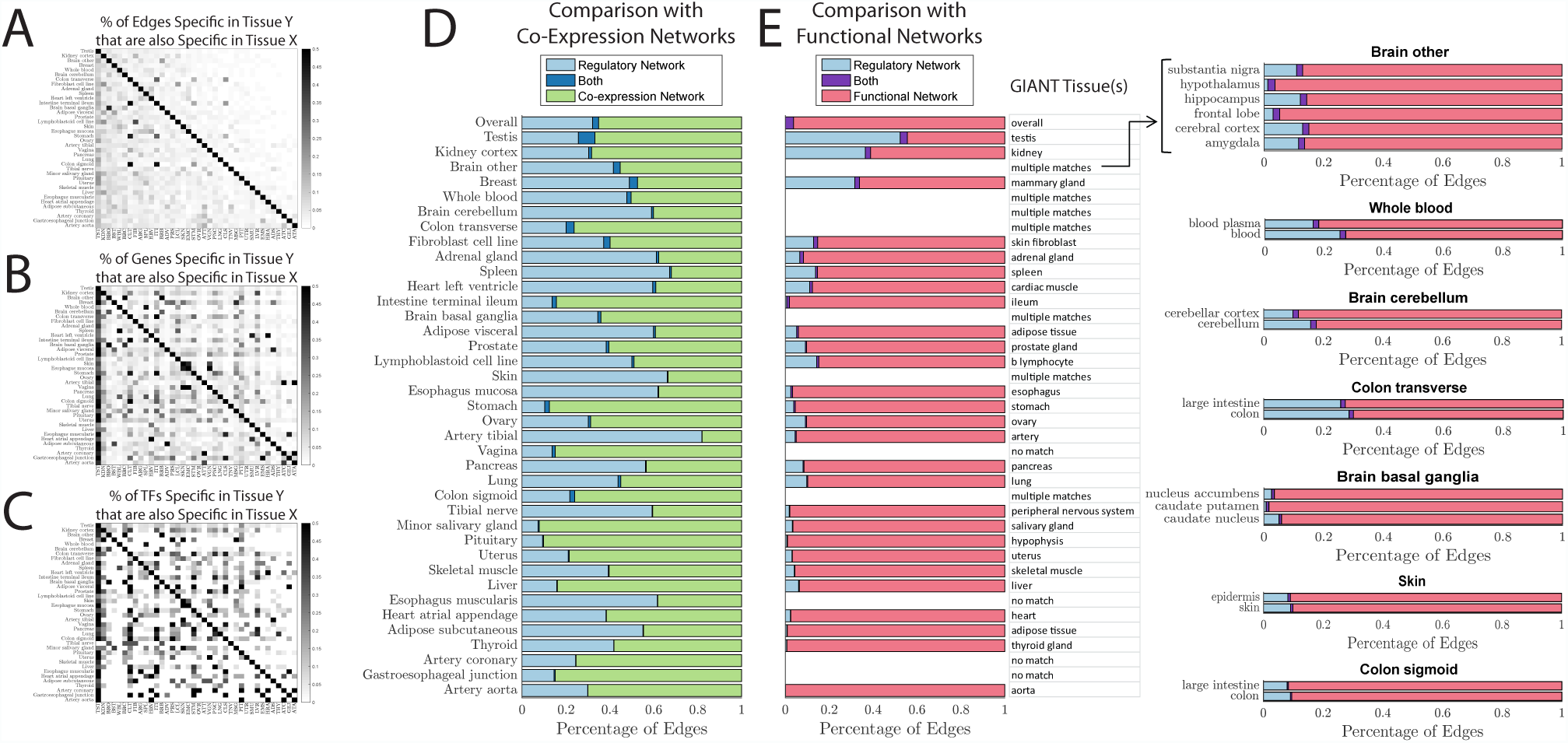
Percentage of (A) edges, (B) genes, and (C) TFs that were identified as specific in the tissue listed along the Y-axis, that are also identified as specific to the tissue listed along the X-axis. (D) Comparison of the tissue-specific edges identified using PANDA-networks to those that would have been identified using a network defined based on co-expression information. (E) Comparison of the tissue-specific edges identified using PANDA-networks to functional edges identified by GIANT-networks.

#### Identifying Tissue-Specific Transcription Factors

Each of our network models includes information about the targeting profiles of 644 transcription factors (see Section S.2). In analyzing tissue-specific transcription factors (Figure 2 in the main text) we focus on this subset of 644 transcription factors. However, other transcription factors that do not have a corresponding DNA-binding motif are included in our GTEx expression data. To evaluate these transcription factors we compared the genes in our GTEx expression data with a list of 1, 987 genes that encode transcription factors as reported in a previous publication [6]. This list of 1, 987 transcription factor genes includes 1, 798 that have expression information in the normalized GTEx RNA-seq data (see Section S.1) and 1, 795 that are among the 30, 243 target genes used in our network analysis (see Section S.2). Of these 1, 795 TFs, 1, 178 do not have DNA-binding motifs, while 617 have a DNA-binding motif and are part of the set of 644 TFs we used in our regulatory prior. We compared the tissue-specificity and multiplicity levels of the 644 TFs with DNA-binding motifs and the 1, 178 TFs without motifs (Supplemental Figure S3). TFs with a motif were more likely to be identified as tissue-specific than TFs without motifs (34.5% versus 28.5%, Chi-squared test *p* = 9.2 *·* 10^-3^) and also tended to have higher multiplicity levels (Chi-squared test *p* = 6.1 *·* 10^-14^). Multiplicity levels of TF without motifs are comparable to those of all target genes (see Figure 2). Thus, while TFs with DNAbinding motifs are often shared among multiple tissues, TFs without such motifs are more often specific to only a single tissue, and appear to behave, at least in terms of their expression levels, more like non-TF genes. We note that although these differences may reflect biases in motif databases, they also potentially indicate that there are particular chemical and/or biological properties that have allowed certain transcription factors to be associated with known DNA-binding sequences.

### S.5. Comparison of PANDA and Correlation-Based Networks

Since co-expression networks have been widely used to analyze gene expression data, including in another network analysis of tissue-specificity in GTEx [7], we compared the tissue-specific edges defined based on PANDA-networks to those defined based on co-expression. For each of the 38 GTEx tissues analyzed we created coexpression networks by calculating the Pearson correlation between the 644 TFs and 30, 243 genes included in our network model (see Section S.2). We identified tissue-specific edges in these correlation-based networks using same protocol we used for genes and PANDA edges (Equation S2, with *N* = 2). When we compared the edges identified as tissue-specific using the correlation-based networks to those identified based on the PANDA-reconstructed regulatory networks we found very little overlap (Supplemental Figure S2D).

**Supplemental Figure S3:**
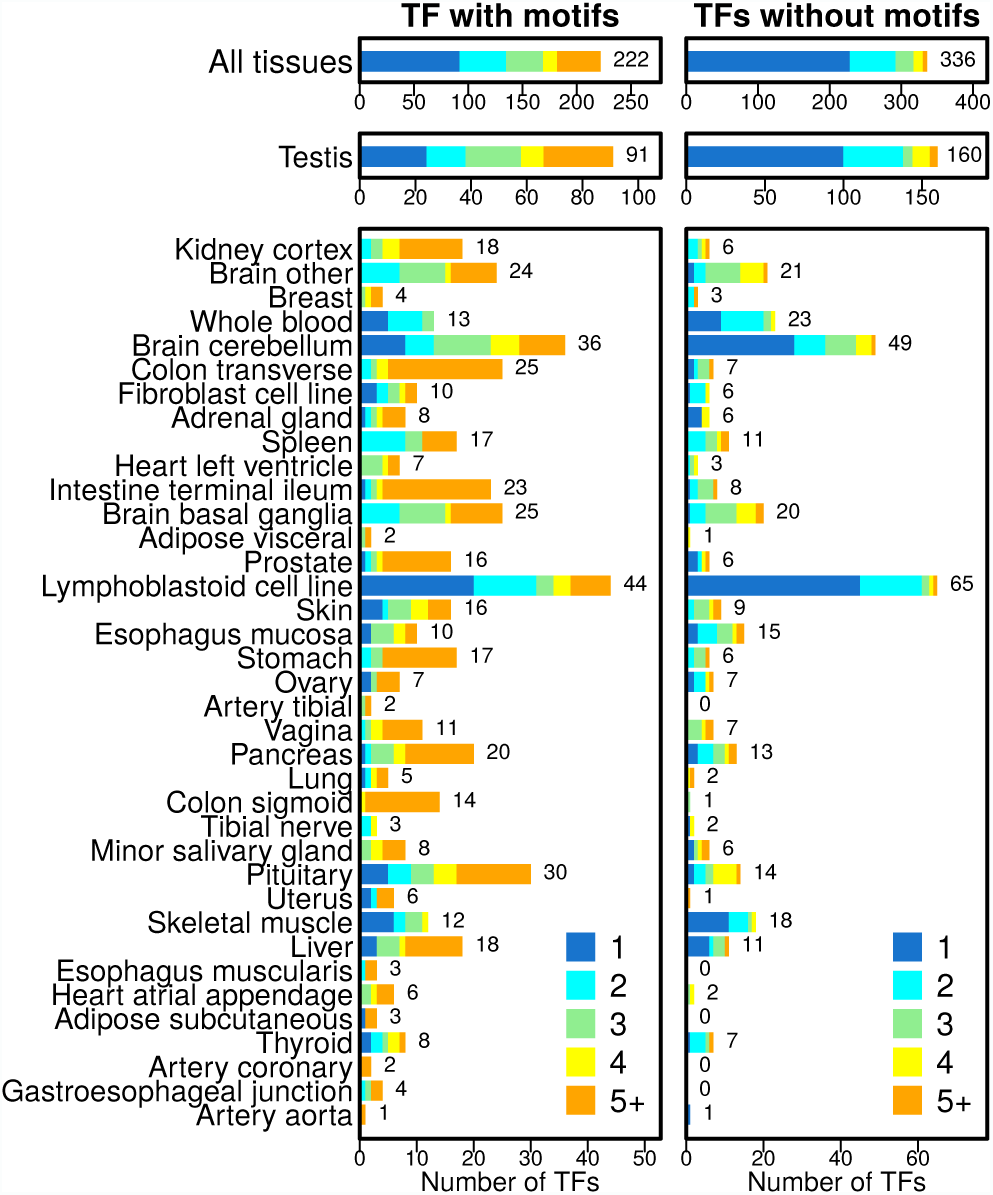
Bar plots illustrating the number of TFs with (A) and without (B) DNA-binding motifs that were identified as “specific” to each of the 38 GTEx tissues. The total number of tissue-specific TFs identified for each tissue is shown to the right of each bar. Tissues are ordered on the number of edges specific to each tissue, as in Figure 2. Tissue-specificity for TFs was defined based on a TF having increased expression in one tissue compared to others, thus some TFs were identified as specific to multiple tissues. This multiplicity value is indicated by the color of the bars. We found that TFs with DNA-binding motifs have substantially higher multiplicity levels compared to TFs without motifs.

This low level of overlap means that PANDA and Pearson correlation networks capture fundamentally different aspects of each tissue’s gene expression program. The co-expression networks are based on measured expression correlations between TFs and their targets. In contrast, PANDA uses co-expression between target genes, not TFs and their targets. In particular, PANDA integrates target co-expression information with a prior regulatory network structure and TF-TF protein-protein interaction data, iteratively updating the likelihood of interactions between TFs and target genes based on shared patterns across these data.

We believe that PANDA more accurately captures tissue-specific regulatory processes. Indeed, when developing PANDA, we compared it to other methods, including co-expression networks, and found that the PANDA networks were better supported by confirmatory data, such as ChIP experiments [4]. Although no ChIP data are available for GTEx, PANDA does find biologically relevant associations that help elucidate the link between expression and tissue phenotype.

### S.6. Comparison of PANDA and Functional Networks

We also systematically explored what, if any, information is shared between the PANDA tissue-specific regulatory networks we estimated from the GTEx data and other published tissue-specific functional networks [8]. In particular, we compared our networks to those estimated by GIANT (Genome-scale Integrated Analysis of gene Networks in Tissues; http://giant.princeton.edu/). This resource aims to capture tissue-specific functional interactions by using tissue-specific knowledge from the literature to selectively upweight particular datasets within a compendium of gene expression information and perform a tissue-ontology aware regularized Bayesian integration.

To begin, we identified which of the 144 distinct tissue-networks available on the GIANT web-resource are most likely to correspond to each of the 38 tissues we analyzed in the GTEx data. In some cases, we identified multiple tissue-networks available from GIANT to compare with a single tissue-network in our GTEx analysis. For example, in constructing the GTEx networks we combined gene expression samples from anatomically close regions of the brain when those samples were indistinguishable using principal component analysis (see Section S.1 and [3]).

Next, we downloaded the top edges identified by GIANT, which represent interactions with a posterior probability greater than 0.1 in that tissue, for each of these identified corresponding tissues. As with the co-expression networks analyzed in Section S.5, these GIANT networks are undirected, with edges extending between pairs of genes. Therefore, we next matched the nodes and dimensions between these functional networks and the GTEx regulatory networks. Specifically, of the 25, 824 Entrez genes found across the GIANT networks, we identified 18, 431 that were uniquely associated with the Ensembl and Gene Symbol annotations for the 30, 243 genes that we had used to reconstruct the GTEx regulatory networks. This represents 91% of the unique Entrez gene annotations in the GTEx data (10, 122 of the GTEx genes did not have a corresponding Entrez annotation). This set of 18, 431 genes included 635 of our 644 transcription factors. We subsetted both the downloaded GIANT networks and our GTEx networks to only consider tissue-specific edges that extend from one of these 635 TFs to one of these 18, 431 genes.

Finally, we determined the number of distinct and common edges between these subsetted GIANT and GTEx networks. We note that the number of top edges in the downloaded GIANT tissue-networks tended to be much greater than the number of tissue-specific edges we had identified from the GTEx PANDA networks. Even so, we found very little overlap between these functional networks and the gene regulatory networks (Supplemental Figure S2E), highlighting that the regulatory information contained in our GTEx regulatory networks is distinct from the type of information that is included in other tissue-network resources.

### S.7 Comparison with a Previously Published Tissue-Specific TF Resource

We also compared the transcription factors we identified as tissue-specific based on the GTEx expression data (see Section S.4) with those reported as tissue-specific in a previous publication [6] (hereafter referred to as NRG, standing for the journal in which it was published: Nature Reviews Genetics) and which were used in other GTEx network evaluations [7]. The results of this analysis are shown in Supplemental Figure S4.

To begin, we downloaded the gene expression data used for the calling of tissue-specific transcription factors in the NRG publication from the Gene Expression Omnibus (GSE1133). We RMA-normalized these expression data using the justRMA() function in the affy Version 1.52.0 library from Bioconductor in R and used a custom-CDF for the Affymetrix GeneChip HG-U133A array (hgu133ahsensgcdf 20.0.0) [9] in order to normalize with respect to current Ensembl genes IDs. This RMA-normalized version of the expression data contained expression information for 11, 900 different Ensembl genes across 158 total samples, 64 of which correspond to the “32 healthy major tissues and organs” used in the NRG analysis. 11, 363 of the genes in this RMA-normalized NRG expression data set also appeared in the normalized GTEx data (see Section S.1 and Supplemental Figure S4A).

We next downloaded the supplemental data that accompanied the NRG manuscript. The “supplemental information S3” file contained information for 1, 987 genes that encode transcription factors, including their “Ensembl gene IDs (release 51), HGNC identifiers, IPI IDs, associated DNA-binding Interpro domains and families, and tissue specificity if any.” Of the 1, 987 transcription factors in this supplemental data file, 1, 130 were included in the RMA-normalized expression data we had downloaded from GEO and 1, 798 had expression information in the normalized GTEx data.

1, 120 of these transcription factors had gene expression values in both the RMA-normalized NRG data and the normalized GTEx data (Supplemental Figure S4A). We evaluated how many of these transcription factors had the same tissue-specific designation in both the NRG supplemental data file and based on our analysis (see Section S.4). To do this we created a map between the 38 tissues used in our current GTEx analysis with the 32 tissues analyzed in the NRG paper. In several cases multiple different GTEx tissue subregions (eg the atrial appendage and left ventricle of the heart) were mapped to the same, more general tissue-designation in the NRG data (eg “heart”). We then directly compared the set of transcription factors that were identified as specific to a given tissue in our GTEx analysis, with the set of transcription factors that were identified as specific to that tissue in the NRG analysis.

**Supplemental Figure S4:**
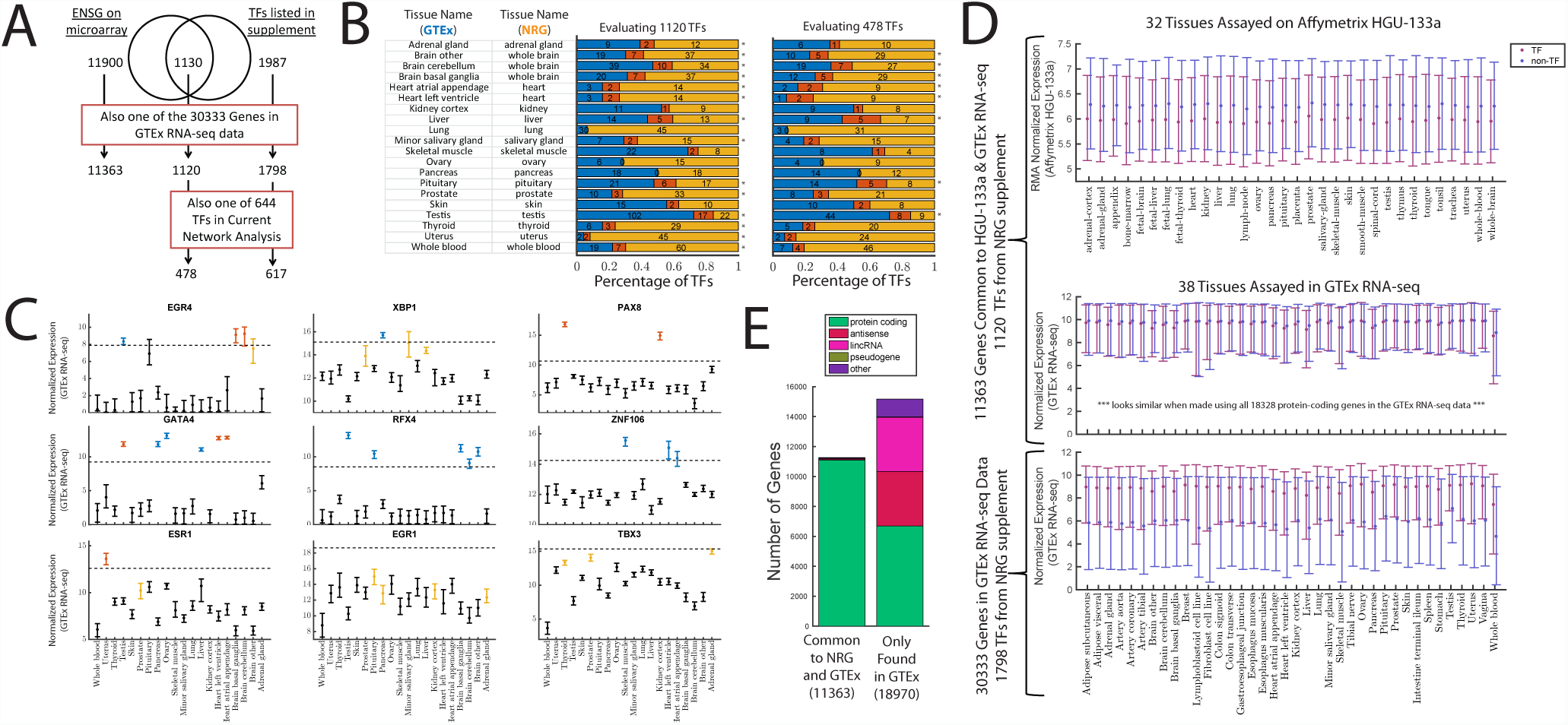
Analysis comparing the results of from a previous publication (NRG) with those obtained in this analysis using the GTEx RNA-seq data. (A) An overview of the overlap in the genes included in the NRG gene expression data, the TFs included in the NRG supplemental data file, and how those sets overlap with the 30, 333 genes in the normalized RNA-seq data we used in this analysis (see Section S.1). (B) An analysis comparing the overlap of TFs identified as specific based on the NRG publication and those identified based on the GTEx data (see Section S.4). An asterisk () indicates that the overlap is nominally significant (*p* < 0.01 by Fisher’s exact test). (C) The distribution of expression values in the GTEx data for several example TFs. These TFs were chosen to illustrate a range of possibilities, including some overlap (*EGR4*, *GATA4*, *ESR1*), as well as opposing (*XBP1*), identical (*PAX8*), or distinct (*RFX4*, *ZNF106*, *EGR1*, *TBX3*) tissue-specific calls based on using either the NRG or the GTEx analysis. As there was little overlap between NRG and GTEx, the four plots with distinct tissue-specific calls are the most representative. (D) The expression of transcription factors versus non-transcription factor genes in both the NRG and GTEx expression data and using various criteria. (E) Information regarding the types of genes that are common between the set on the NRG microarray and in the GTEx RNA-seq data, and the types of genes that we have included in our GTEx expression analysis that were not on the NRG microarray.

We find that the overlap between these sets of TFs is nominally statistically significant in most cases (*p* < 0.01 in 14 of the 20 comparisons). However, the actual number of TFs identified as specific to a particular tissue in both the NRG and our GTEx analysis is quite low (Supplemental Figure S4B). For example, the lung, ovary, and pancreas contained no common tissue-specific TFs between our GTEx designation and the NRG-designation. In addition, when we restrict this analysis to the 478 of these 1, 120 TFs that were also included as regulators in our network model, even this nominal significance goes away for many tissues.

To better understand this result, we examined the distribution of expression values in the GTEx data for these 1, 120 TFs. A few examples are included in Supplemental Figure S4C. In some cases, such as for *XBP1* and *TBX3*, the fact that a TF was only identified as specific by NRG and not GTEx appears to be a function of the cutoff we used for defining tissue-specificity (see Section S.4). However, we note that relaxing this criterion would have significantly changed the number of TFs we identified as tissue-specific (see Supplemental Figure S1B) and ultimately would not affect the relatively low level of overlap we see here. In addition, there are many examples where our GTEx analysis clearly identifies tissue-specific signals that are not reflected in the NRG data set (*ZNF106*, *RFX4*, *GATA4*), and also examples where there is no apparent tissue-specific signal for a TF despite it being called so in the NRG data (*EGR1*, *ESR1*). Given that the NRG expression data contains only two samples per tissue, we believe that the tissue-specificity calls for TFs made in our analysis are more reliable.

This low level of overlap in the identified tissue-specific TFs led us to more closely investigate the expression data used in the NRG analysis. Using the RMA-normalized NRG data (and focusing on the 11, 363 genes and 1, 120 TFs that are common between the NRG and GTEx data sets), we reproduced the plots from Figure 3 in the NRG publication. Consistent with that analysis, we find that in the NRG expression data set transcription factors are expressed at lower levels than non-TFs (compare Figure 3A in [6]) to Supplemental Figure S4D). We then repeated this same analysis using the GTEx data. To our surprise, the difference in expression between TFs and non-TFs largely disappeared when performing this analysis in the GTEx data. Finally, we repeated this analysis using all 30, 333 genes in our GTEx expression data set. This actually resulted in the opposite conclusion as the analysis presented in the NRG paper, with TFs expressed at higher levels than non-TFs.

One advantage of using RNA-sequencing data over microarrays is that sequencing can capture mRNA from many different types of genes and is not limited by the set of probes included on a given array. To better understand whether differences in technology (microarray versus RNA-sequencing) may be influencing the results shown in Supplemental Figure S4D, we determined the annotations for the 30, 333 genes included in our GTEx analysis using Biomart (dec2013.archive.ensembl.org). Supplemental Figure S4E shows the distribution of these annotations across the 11, 363 genes that are common between the NRG microarray and the GTEx RNA-seq data, and across the 18, 970 genes that are only contained in our GTEx RNA-seq data. It is immediately clear that the microarray genes are almost completely composed of protein-coding genes whereas the genes captured only in the GTEx data contain many types, including antisense, lincRNAs, and pseudogenes. Thus the fact that we see TFs expressed at higher levels than non-TFs when evaluating the full 30, 333 genes in the GTEx data is largely a consequence of the fact that all TFs are, by definition, protein-coding genes, and that protein-coding genes are expressed at higher levels than non-protein-coding genes.

Overall, this analysis highlights the importance of the public availability of data and reproducible research, as we were able to faithfully reproduce many of the results from the NRG paper using their original data. It also highlights the need to revisit previous analyses as new data becomes available. The differences in tissuespecificity and TF-expression based on the NRG analysis and the GTEx data are a perfect demonstration of the opportunity the GTEx data gives us to revisit our understanding of tissue-specificity and gene regulation.

### S.8. Calculating Enrichment of Tissue-Specific Edges

To quantify the relationship between various tissuespecific edges and nodes, we explicitly evaluated the extent to which tissue-specific edges are more (or less) likely to target tissue-specific genes (or TFs) as compared to chance. For each of the 38 tissues we counted the number of edges called as specific to a tissue (*t*, see Equation S2), and of a given multiplicity (*M*, see Equation S3) that also target a gene identified as specific to that tissue (see Equation S4):

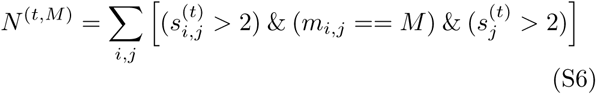

We then summed these numbers over all 38 tissues:

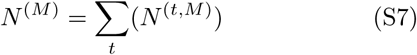

We also calculated the number of tissue-specific edges of a given multiplicity that one would expect to target tissue-specific genes by chance:

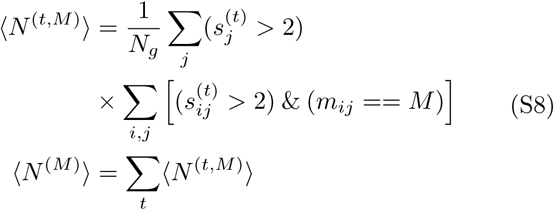

Where *N*_*g*_ = 30, 243 (the number of genes in our model). Finally, we defined the enrichment for tissue-specific edges of a given multiplicity targeting tissue-specific genes as:

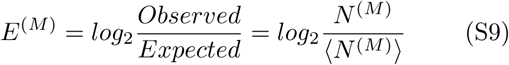

We found very high enrichment for tissue-specific edges targeting tissue-specific genes, especially in edges with lower multiplicity values (Figure 3A).

### S.9. Gene Set Enrichment on TF Targeting Profiles

Gene Set Enrichment Analysis to Quantify the Functions Associated with Tissue-Specific TF-targeting: Although tissue-specific transcription factors are more likely to be associated with tissue-specific network edges than one would expect by chance, we found that this association is much lower than the association between tissue-specific edges and target genes. This led us to the hypothesis that both tissue-specific and non-tissue-specific transcription factors play an important role in mediating tissue-specific biological processes. To test this hypothesis, for each transcription factor (*i*), we quantified its tissue-specific targeting profile in a given tissue (*t*) as 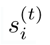 (see Equation S2). We then ran a pre-ranked Gene Set Enrichment Analysis (GSEA) [10] on the scores in this profile to test for enrichment for Gene Ontology (GO) terms. In total we performed 24, 472 GSEA analyses, one for each of the 644 transcription factors included in the network for each of the 38 tissues. The detailed results of this analysis for each tissue are given in Supplemental Tables 4 and 5.

#### Selection of TFs with Highest and Lowest Expression Enrichment

In order to better understand the relationship between tissue-specific transcription factor expression patterns and their tissue-specific targeting of biological functions, we selected ten transcription factors with the highest expression enrichment based on Equation S4. More specifically, for the analysis presented in Figure 4B in the main text, we selected the ten transcription factors with the highest 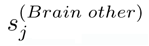 value, and the ten transcription factors for which the absolute value of 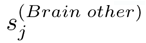 was closest to zero.

#### Identifying Differentially Targeted Biological Processes and Differentially Targeting TFs for Each Tissue

For each tissue, we identified GO terms that were significantly enriched (*F DR* < 0.001; GSEA Enrichment Score, *ES >* 0.65) for tissue-specific targeting by at least one transcription factor. This allowed us to define 38 sets of differentially targeted biological processes, one for each tissue. For each tissue, we used the corresponding set of differentially targeted GO terms to identify differentially targeting TFs. More specifically, for each tissue we determined the set of TFs that were specifically significantly-enriched (*F DR* < 0.001; GSEA Enrichment Score, *ES >* 0.65) for differential targeting of at least one of the members in the complete set of differentially targeted biological processes. This allowed us to define 38 sets of differentially targeting TFs, one for each tissue. Interestingly, these TFs were not associated with the sets of differentially expressed (tissue-specific) TFs identified in Section S.4 (Supplemental Figure S5 and Supplemental Table 5).

#### Community Structure Analysis to Identify Related Sets of TFs/Tissues and GO terms

To gain a more holistic understanding of the patterns of tissue-specific targeting across all 38 tissues, we combined the GSEA analysis results into a single large matrix that contained the enrichment results across all 24, 472 transcription factor and tissue pairs. This matrix contained all the tested GO terms in the rows, and each of the 24, 472 GSEA analyses in the columns. We selected elements of this matrix that represented highly significant positive enrichment for tissue-specific targeting (*F DR* < 0.001 and *ES >* 0.65), creating a bipartite network where nodes were either GO terms or TF-tissue pairs (the pairs used for the GSEA analyses). We then ran the fast greedy community structure detection algorithm [11] to identify “communities,” or sets of GO terms associated with TF-tissue pairs, in this bipartite network. The benefit of this type of analysis over other clustering approaches, such as hierarchical clustering, is that each “node” is assigned to exactly one community, aiding in our interpretation of these highly complex results. This analysis identified 48 separate communities (Figure 5A and Supplemental Figure S6), or clusters of GO terms associated with TF-tissue pairs (representing the tissue-specific targeting profile of a particular TF in a particular tissue). All TF-tissue pairs with significant positive enrichment tissue-specific targeting of a particular GO term can be found in Supplemental Table 6. Characteristics of the 48 communities found by clustering these relationships can be found in Supplemental Table 7.

**Supplemental Figure S5:**
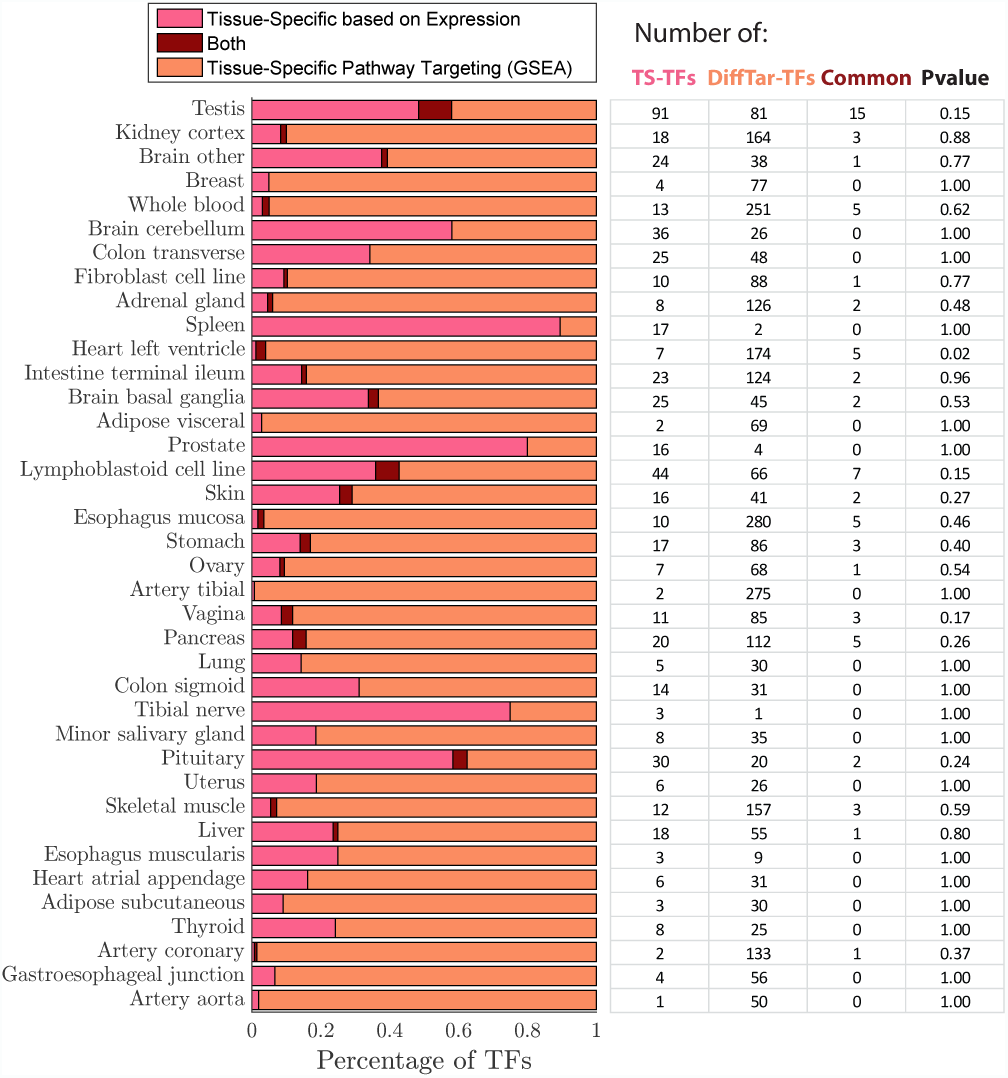
Comparison of TFs defined as tissue-specific based on their expression profile, versus based on their differential targeting profile. All TFs that have tissue-specific differential targeting profiles can be found in Supplemental Table 5.

#### Word Clouds to Visualize the Functional Content of Communities

Nine communities had eight or more GO term members. For these communities we summarized their functional content using a free word cloud making program (downloaded from: http://www.softpedia.com/get/Office-tools/Other-Office-Tools/IBM-Word-Cloud-Generator.shtml). This program automatically configures the orientation of words in the clouds, but we manually assigned each word a relative size based on that word’s statistical enrichment in the community [12]. Specifically, for a given community, we counted the number of times an individual word appeared across all the GO term members associated with that community (Nwc) and then calculated its statistical enrichment in a given community based on the hypergeometric probability:

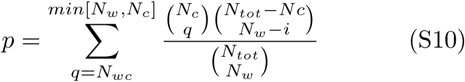

where *N*_*c*_ is the number of individual words in a community, *N*_*w*_ is the number of times the word appears across all term descriptions and *N*_*tot*_ is the total number of words included in all tested GO terms. We then scaled the sizes of the words in the word cloud based on –*log*_10_(*p*) such that words that have the lowest probability of being in the community by chance are given the largest size and words that are common across many biological functions and that one might expect to be in a community by chance are given a very small size.

**Supplemental Figure S6:**
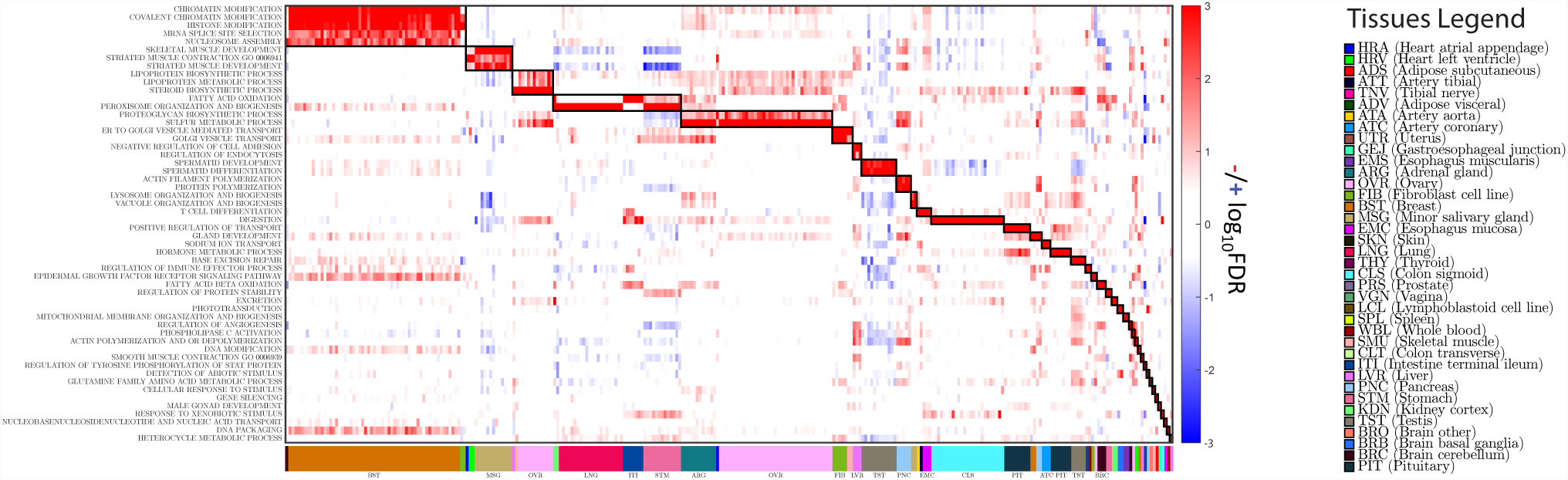
Illustration of the communities of GO terms and TF-tissue pairs that had three or fewer GO-term members.

#### Bipartite Network to Visualize the Relationships between Communities and Tissues

Figure 5C was made using JavaScript library D3.js (http://bl.ocks.org/NPashaP/fcb09e2cddbe104e209f457d44f166ca).

#### Transcription Factor Enrichment in Communities

Because of the complex structure of the relationships represented between TF/Tissue pairs and the GO Terms in our communities, we identified transcription factors that were significantly enriched in a given functional community by performing a permutation analysis. We began by determining the number of times each transcription factor has a significant GSEA association with each of our 48 functional communities. Then, to determine whether this value was greater than expected by chance, we performed a supervised shuffling of the community labels of the transcription factors. In particular, to perform a community label shuffling that would correctly identify enrichment among transcription factors, we first identified the set of community assignments associated with each tissue, and shuffled these assignments only among the TF/Tissue-pairs for that tissue. This approach allowed us to conserve both the size of the communities and the distribution of tissues within the communities. After performing each shuffling, we counted the number of times each TF had an association with each community in this random assignment. We repeated this shuffling 10, 000 times and estimated the significance of enrichment of a transcription factor in each community by determining the percentage of times the counts from the shuffled assignments were greater than the counts from the original assignments.

### S.10. Network Centrality Estimates of Tissue-Specific Genes

We used the igraph Version 1.0.0 package in R to calculate both the degree (using the graph.strength() function) and betweenness centrality (using the between-ness() function) of genes in each of the 38 complete, weighted PANDA tissue networks (see Section S.2 and Equation S1).

Degree: The degree of a node is defined as the number of edges connected to that node. Because we have weighted graphs, we calculated the degree of a gene in a given tissue (*t*) by summing up the weights of all edges connected to that gene (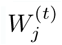 see Equation S1). Note that because these are also complete graphs, each gene had exactly 644 edges, one from each transcription factor.

Betweenness: The betweenness of a node is defined as the fraction of non-redundant shortest paths in the network that go through that node. In a weighted network, the shortest path calculation uses edge weights to calculate the cost of traversing each edge. In order to prefer higher edge weights in calculating shortest paths, we used 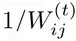 (see Equation S1) as the cost for determining the shortest paths. In order to calculate the betweenness centrality, we treated edges as undirected (meaning that an edge exists both from a TF to its target gene and from the target gene to the TF).

### S.11. Network Centrality of PANDA’s Seed Regulatory Network

PANDA builds its predicted regulatory network, in part, by leveraging information from a prior “seed” network constructed by mapping transcription factors to genes based on genome sequence information (see Section S.2). We wanted test whether the differences in centrality values that we observed between tissue-specific and non-tissue-specific genes were due to the structure of this input data or if they were identified primarily through PANDA’s message passing network optimization. Therefore, we calculated the degree and betweenness centrality for genes based on the motif scan seed network (see Section S.10). We note that this seed network is “unweighted,” meaning that the edges only take two values: one if the motif for TF *i* is found in the promoter region of gene *j*, and zero if it is not.

In the motif prior network, we saw only minimal differences between the centrality of tissue-specific and non-tissue-specific genes, with tissue-specific genes having slightly lower centrality values compared to non-tissue-specific genes (Supplemental Figure S7). This is consistent with our finding in the main text that tissue-specific genes are generally of low betweenness and only see an increase in their betweenness in their “specific” tissues, and supports our interpretation that tissue specificity is associated with increased centrality in the network as genes gain new non-canonical regulatory paths.

**Supplemental Figure S7:**
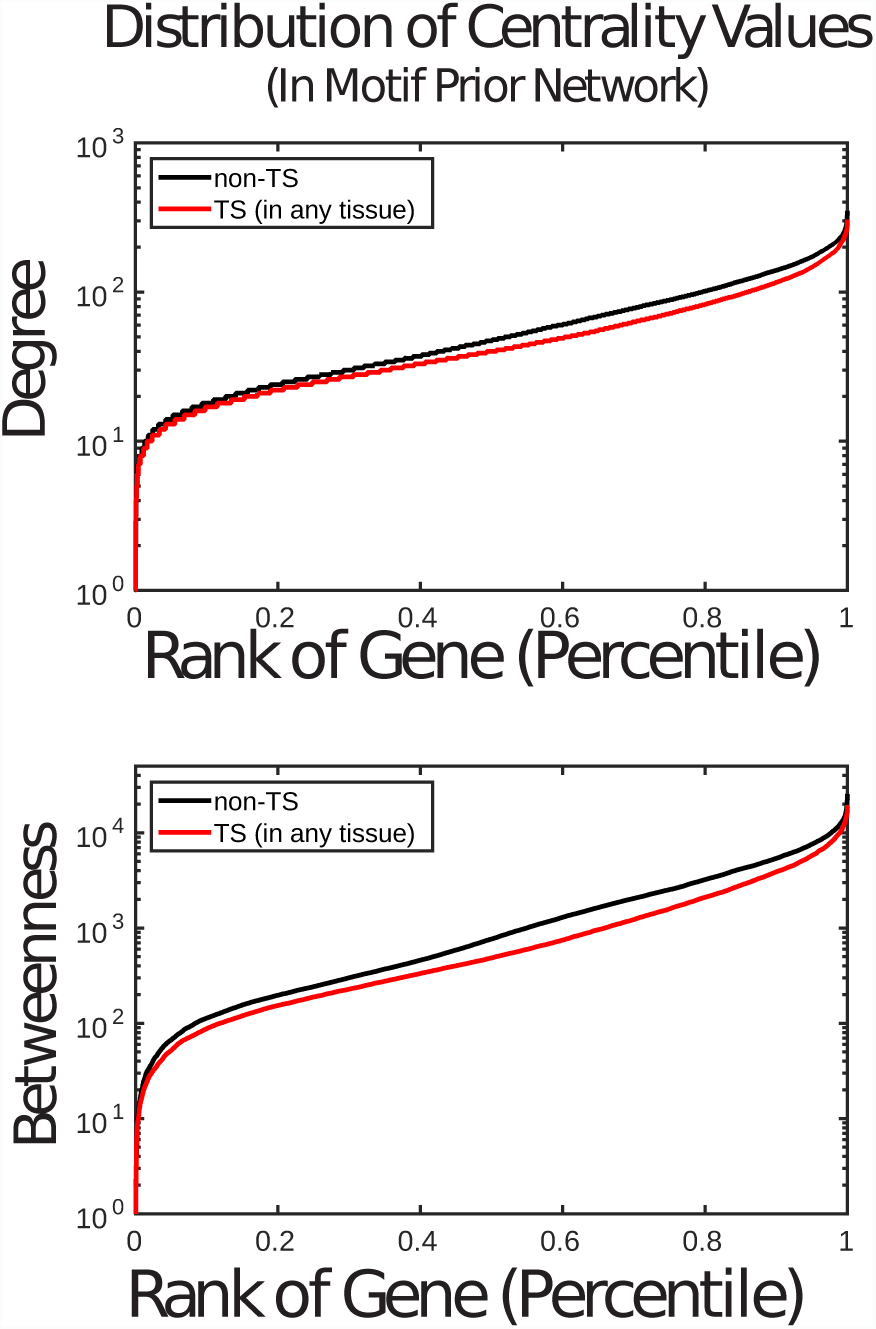
Distribution of the (A) in-degree and (B) betweenness centrality values of genes in the motif prior network used to seed the PANDA algorithm. Genes identified as tissue-specific are represented in the red line (all multiplicities considered), while those that are not identified as specific to any tissue are represented by the black line.

### S.12. S-12 Identifying Genes with eQTLs and GWAS Variant Associations

To evaluate the potential role of genetic variants in tissue-specific gene regulation, we identified tissue-specific, *cis*-acting expression quantitative trait loci (eQTLs), as described in [13]. Briefly, of the 38 tissues for which we had reconstructed gene regulatory network, 19 contained gene expression samples from at least 150 distinct individuals with imputed genetic data (note that only tissues with at least 200 individuals were presented in [13], but the data were processed the same way). These 19 tissues included adipose subcutaneous, artery aorta, artery tibial, brain other, breast, fibroblast cell line, colon transverse, esophagus mucosa, esophagus muscularis, heart atrial appendage, heart left ventricle, lung, skeletal muscle, tibial nerve, pancreas, skin, stomach, thyroid, and whole blood. For each of these tissues, we identified single-nucleotide polymorphisms (SNPs) that had a minor allele frequency greater than 0.05 across the individuals with associated tissue-specific gene expression data and used Matrix eQTL [14] to quantify the statistical association of the expression of each of the 29, 155 genes in GTEx with each of these genetic variants. For this analysis we used a *cis*-acting window around the gene of 1 mega-base, and adjusted for sex, age and the three first principal components obtained using the genotyping data. Finally, we determined which genes had at least one significant (*F DR* < 0.05) eQTL association in each tissue.

We also determined which of these QTL-associated genes might be important for disease or other phenotypic traits. To do this, we downloaded the NHGRI-EBI GWAS Catalog (http://www.ebi.ac.uk/gwas/; access date: 12/08/2015). We curated this information, excluding any entries in the catalog for which the variant did not have an associated rsid. Then, we parsed our identified tissue-specific eQTL associations, pruning those that were not with one of these GWAS genetic variants. Finally, we used this information to determine which genes had at least one significant (*F DR* < 0.05) eQTL association with a GWAS-SNP in each tissue. These genes are listed in Supplemental Table 8.

We compared the genes identified in these analyses with information regarding tissue-specificity. In particular, since we only identified eQTL associations in 19 tissues, we identified a new set of “tissue-specific genes”, which included those that had a multiplicity greater than zero when applying Equation S5 and summing only over the 19 tissues. 5, 256 of the 29, 155 genes were identified as specific to at least one of these 19 tissues.

## SUPPLEMENTAL TABLE LEGENDS

Supplemental tables are available online.

Supplemental Table 1: Table listing edges that are either scaled the sizes of the words in the word cloud based on in the regulatory network prior, or are identified as tissuespecific.

Supplemental Table 2: Table listing the genes included in our PANDA network models, including their multiplicity, tissue-specificity based on gene expression information, and centrality values.

Supplemental Table 3: Table listing the transcription factors included in our PANDA network models, including their multiplicity, tissue-specificity based on gene expression information, and centrality values.

Supplemental Table 4: Table listing the percentage of genes and TFs associated with a tissue-specific edge in each tissue.

Supplemental Table 5: Table listing each of the 38 tissues included in our analysis, the GO terms identified as having significantly increased targeting in each tissue (*F DR* < 0.001 and *ES >* 0.65 by at least one transcription factor) and the TFs that are differentially targeting these categories.

Supplemental Table 6: Table listing all significant GSEA results (*F DR* < 0.001 and *ES >* 0.65) obtained in our differential targeting analysis.

Supplemental Table 7: Table listing statistics for the 48 “communities” of GO-terms and TF-tissue pairs that were identified when clustering the GSEA results. The “top category” is the GO term with the largest number of significantly associated TF-tissue pairs. The table also includes all the GO-terms and TF-tissue-pair members in each of the 48 “communities”.

Supplemental Table 8: Table listing the 308 genes that had a tissue-specific eQTL with one of the genetic variants listed in the GWAS catalog. The tissue in which these eQTLs were identified, the tissue(s) where the associated gene was identified as specific (among the 19 in common between the eQTL and network analysis), and any common tissues are also listed.

## SUPPLEMENTAL FILES

Supplemental File 1: Interactive version of Figure 5C.

